# Human Gastric Multi-Regional Assembloids Favour Functional Parietal Maturation and Allow Modelling of Antral Foveolar Hyperplasia

**DOI:** 10.1101/2024.07.08.602480

**Authors:** Brendan C Jones, Giada Benedetti, Giuseppe Calà, Lucinda Tullie, Ian C Simcock, Roberto Lutman, Monika Balys, Ramin Amiri, Jahangir Sufi, Owen Arthurs, Simon Eaton, Glenn Anderson, Nicola Elvassore, Vivian SW Li, Kelsey DJ Jones, Christopher J. Tape, Camilla Luni, Giovanni Giuseppe Giobbe, Paolo De Coppi

## Abstract

Patient-derived human organoids have the remarkable capacity to self-organise into more complex structures. However, to what extent gastric organoids can recapitulate human stomach physiological functions remain unexplored. Here, we report how region-specific gastric organoids can self-assemble into complex multi-regional assembloids showing functional response to drugs targeting the ATPase H+/K+ pump. The assembloids show preserved fundus, body, and antrum regional identity, and gastric-specific crosstalk pathways arise. The increased complexity and cross-communication between the different gastric regions, allow for the emergence of the elusive parietal cell type, responsible for the production of gastric acid, with functional response to drugs targeting the ATPase H+/K+ pump. Remarkably, we generated assembloids from PMM2-HIPKD-IBD paediatric patients (Phosphomannomutase 2 – Hyperinsulinemic hypoglycaemia and autosomal recessive polycystic kidney disease - Inflammatory bowel disease), a genetic condition found to be associated with unusual antral foveolar hyperplasia and hyperplastic polyposis. The cellular mechanisms behind such phenomena are poorly understood, and an exhaustive experimental model is needed. The ΔPMM2 multi-regional assembloid we have generated efficiently recapitulates hyperplastic-like antral regions, with decreased mucin secretion and glycosylated ATP4b, which results in impaired gastric acid secretion. Multi-regional gastric assembloids, generated using adult-stem cell-derived organoids, successfully recapitulate the structural and functional characteristics of the human stomach, offering a promising tool for studying gastric epithelial interactions and disease mechanisms previously challenging to investigate in primary models.

## Introduction

The human stomach is a vital organ responsible for the initial phases of food digestion that relies upon finely regulated hormonal functions. As a complex and dynamic system, the stomach plays a crucial role in the overall digestive process, enabling the breakdown of ingested food into absorbable nutrients. Structurally, it consists of three main distinct regions: fundus, body, and antrum, each with contrasting morphological and functional properties^1,2^. Two types of gastric glands with distinct locations have been described: oxyntic and antral glands. The oxyntic glands, located in the fundus and body, are characterized by an abundance of parietal cells and mucin-secreting cells in the neck, and are essential for the secretion of hydrochloric acid (HCl) and intrinsic factor (IF). The antral glands have a longer pit domain with neck mucous and endocrine cells, predominantly G cells at the base, and are responsible for producing the gastrin hormone, which plays a central role in regulating gastric acid secretion in the fundus^2,3^.

During human development, the antrum forms before the fundus, usually around the fourth week of gestation. The antrum develops from the caudal portion of the primary stomach, and later the fundus develops from the cranial portion^4^, with regional specification occurring in the early stages of embryonic formation. Animal models are not suited to investigate these interactions, i.e., the mouse stomach consists of corpus and forestomach only, the latter presenting a keratinizing stratified epithelium^5^. In the last decade, primary human organoid models, which consist of gastric epithelial cells only, have emerged as useful tool for modelling various physiological and pathological processes in the stomach, thanks to their ability to recapitulate certain functions, retain genetic diversity and patient-specific features^6–10^. Alternatively, gastric organoids derived from human pluripotent stem cells (hPSC) are advantageous as they can also contain mesenchyme. However, in contrast to primary organoids, achieving regionalization in hPSC-derived organoids is more complex and lengthier using current protocols^10^.

While primary organoids provide valuable insights into gastric mechanisms, they are still relatively simple models when compared to native tissue, lacking features that are challenging to reproduce *in vitro*. Similar limitations were observed in organoids derived from different intestinal regions, which may recapitulate some of the biological characteristics of the specific region, but fail to fully reflect the complexity of epithelial function^9,11^.

One of the major limitations of primary gastric models is related to the absence of more complex cell interactions, which render simple organoids an incomplete tool for modelling the organ of origin^11^. For instance, functional acid-producing parietal cells present in gastric tissue biopsies are poorly represented in murine primary gastric epithelial organoids^12^. On the other hand, examples of murine-derived fundic organoids presenting parietal cells have been reported^13^. Furthermore, the possibility to induce the differentiation of parietal cells has been demonstrated with hPSC-derived fundic gastric organoids^14^. However, the appearance of functional parietal cells has never been reported in human tissue-derived gastric organoids^15,16^, but only in primary non-organoid models^17^.

Investigating the dynamic interplay between the antrum, body, and fundus is essential for enhancing our understanding of the mechanisms that regulate gastric physiological functions, and associated pathologies. For example, disorders related to disrupted acid production within the stomach may be caused by dysregulation of the different regions of the stomach during development or later in life^18^. Gastric acid-related diseases constitute a spectrum of gastrointestinal (GI) conditions that arise from disruptions in stomach functions, exerting profound effects beyond the GI tract. This encompasses ailments such as gastritis, peptic ulcers, and gastroesophageal reflux disease, often stemming from aberrant gastric acid production^18^. To date, treatments have focused on symptoms management, and prevention of long-term complications. However, little has been done to investigate the dysregulated pathology leading to gastric disorders.

To overcome these challenges and advance our understanding of the gastric interactions between the antrum and the fundus, a novel research approach is warranted. In this context, the development of a gastric assembloid model emerges as a promising strategy^19^. The gastric multi-regional assembloids (MRA) we propose here are advanced *in vitro* models that faithfully recapitulate the structural, molecular, and functional characteristics of the human stomach epithelium using patient-derived tissue-specific progenitors.

To assess the applicability of our system for disease modelling, we generated primary gastric epithelial cell organoids from patients with PMM2-HIPKD-IBD (Phosphomannomutase 2-associated Hyperinsulinism with Polycystic Kidney Disease and Inflammatory Bowel Disease). *PMM2* encodes Phophomannomutase 2, which is a non-redundant component in the N-glycosylation pathway. Patients with biallelic deleterious mutations in *PMM2* suffer from a congenital disorder of glycosylation (CDG) that is associated with complex multisystemic manifestations incorporating pervasive neurologic features. In contrast, patients with PMM2-HIPKD-IBD harbour monoallelic deleterious mutation in *PMM2 in trans* with a specific *PMM2* promotor mutation, or the promotor mutation in homozygosity, and have a more restricted pattern of disease incorporating hyperinsulinism (HI), polycystic kidney disease (PKD), inflammatory bowel disease (IBD). These patients also appear to have a tendency towards the development of gastric antral foveolar hyperplasia and antral hyperplastic polyposis in early childhood^20^. Such gastric manifestations are very unusual in children, and have not been identified in association with any other monogenic Inflammatory Bowel Disease syndrome to our knowledge. The cellular and molecular mechanism(s) underlying this tendency is unknown, but in general the organ-level pattern of disease in PMM2-HIPKD-IBD reflects the tissue-specific expression of the transcription factor *HNF4A.* We have hypothesized that the promotor mutation interrupts *HNF4A cis*-acting regulatory control of *PMM2* expression, resulting in critically reduced PMM2 activity in a restricted set of cells and tissues ^20^. Based on published expression data, we have proposed that the gastroenterological manifestations of PMM2-HIPKD-IBD most likely represent an epithelial impairment^20^. Exploiting the potential of the gastric multi-regional assembloids, we were able to model some features of the gastric antral epithelial hyperplasia *in vitro*.

We have demonstrated that the organoids retain native regional specificity after isolation from gastric tissue. When combined to generate MRA, we can generate an *in vitro* system that more closely recapitulates the stomach epithelium. Indeed, we observed the spontaneous emergence of functional parietal cells, that were absent in other epithelial organoid systems in expansion medium. MRA models may enable controlled manipulation, and observation of specific cellular components and signalling pathways, providing insights into spatial interactions, and disease mechanisms that would be otherwise unattainable.

## Results

### Gastric organoids from fundus, body, and antrum retain regional memory

To determine whether gastric stem cells can retain the identity of the region of origin, we isolated mucosal biopsies from the antrum, body, and fundus regions of four paediatric patients (**Fig.1a, Supplementary Table 1**). We verified the presence of the regional markers in paediatric tissues. Antral tissue showed the presence of gastrin (G cells) and PDX1, the latter absent in fundus and body epithelium, while fundus tissue showed defined IRX3 marker expression. (**Fig.1b**, **Extended data Fig.1a**). This is in keeping with PDX1 expression in post-conception week (PCW) 10 human stomach which is limited to the antral cells, showing the early establishment of gastric regionality (**Extended data Fig.1b**)^21,22^.

**Fig.1.**
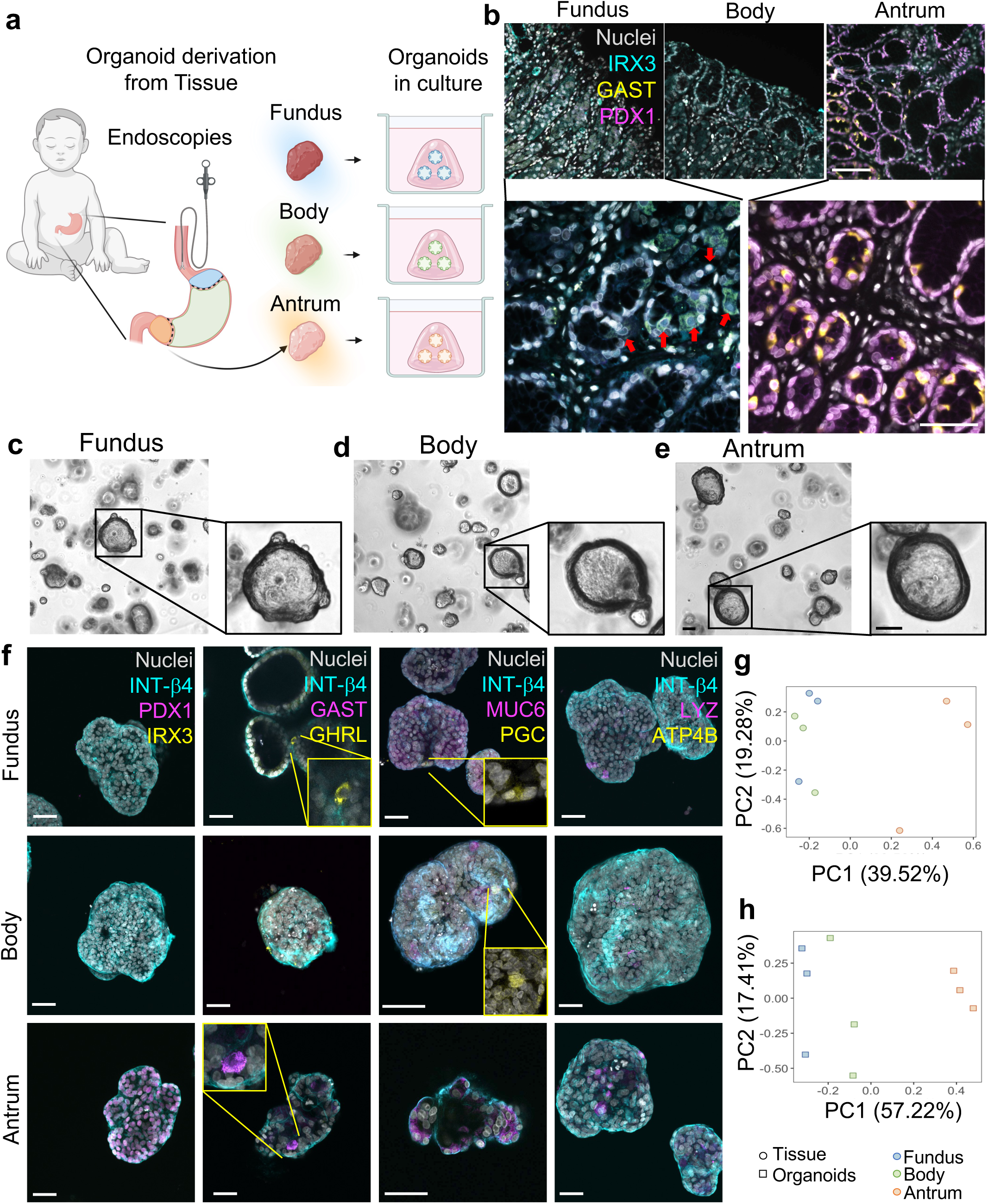
Characterization of gastric regionality in tissues and organoids. **a**, Schematic representation of organoid derivation from paediatric patients’ gastric mucosal biopsies. Created with BioRender.com. **b**, Immunofluorescence panel showing Fundus, Body, and Antrum paediatric stomach tissue sections stained for Iroquois homeobox 5 (IRX5) in cyan, Gastrin (GAST) in yellow, Pancreatic and Duodenal homeobox 1 (PDX1) in magenta, and nuclei in grey (Hoechst). Scale bar 100 μm and 50 μm in the enlargement. **c**, Bright field image of representative organoid line for Fundus region, showing the formation of spherical organoids within 7 days starting from single cells. Scale bar 100 μm. **d**, Bright field image of representative organoid line for Body region, showing the formation of spherical organoids within 7 days starting from single cells. Scale bar 100 μm. **e**, Bright field image of representative organoid line for Antrum region, showing the formation of spherical organoids within 7 days starting from single cells. Scale bar 100 μm. **f**, Immunofluorescence panel showing Integrin-b4 (INT-b4) in cyan; Pancreatic and Duodenal homeobox 1 (PDX1), Gastrin (GAST), Mucin 6 (MUC6), Lysozyme (LYZ) in magenta; Iroquois homeobox 3 (IRX3), Ghrelin (GHRL), Pepsinogen C (PGC), ATPase H+/K+ transporting subunit b (ATP4B) in yellow; nuclei in grey (Hoechst) in Fundus, Body, and Antrum organoids. Scale bar 20 μm. **g**, Principal component analysis (PCA) of RNA-sequencing samples from Fundus (blue), Body (green), and Antrum (red) tissue (circles). (n=3 for biological replicates). **h**, Principal component analysis (PCA) of RNA-sequencing samples from Fundus (blue), Body (green), and Antrum (red) organoids (squares). (n=3 for biological replicates).

Gastric organoids were derived according to previously established protocols^8,23^ and cultured in gastric-specific medium^24^. These organoid cultures generated from stomach fundus, body and antrum showed a comparable cystic morphology (**Fig.1c-e**). Fully grown organoids (day 7 of culture) were immunostained for regional- and gastric-specific markers. Strikingly, PDX1 specifically marked antrum-derived organoids only, in which we detected a low proportion of cells positive to GAST, MUC6, and LYZ. Fundus and body organoids exhibited cells positive to LYZ, MUC6, GHRL and PGC. Importantly, presence of ATP4B was not detected in any of the organoids (**Fig.1f**).

We then characterised native tissues (mucosal controls) and regional organoids with bulk RNA sequencing. Overall, PCA analysis on the tissue showed the antral samples separating from the cluster including fundus and body samples (**Fig.1g**). The same trend was found when analysing organoid samples, with antral organoids clustering separately from the other two regions (**Fig.1h**). When comparing tissue and organoids, we could appreciate the presence of a compact cluster defined by the organoids of all regions. While strongly diverging from the organoid cluster, the tissue samples from the three regions resulted more scattered, with the antrum separating from the other two regions (**Extended data Fig.1c**). Looking at specific regionality targets, *PDX1* was expressed by the antrum in both tissue and organoids. *GAST* (expressed by G-cells in the antrum), *ATP4A* and *ATP4B* (expressed by parietal cells in body and fundus), and *GHRL* (expressed in P/D1 or X cells in body and fundus) were respectively found to be expressed in tissue, whilst almost absent in organoids. *IRX3* (a transcription factor mainly expressed in body and fundus) and *PGC* (expressed in chief cells, more strongly in body and fundus) transcripts were present in tissue, and more sparsely in organoids (**Extended data Fig.1d**).

### Gastric organoids self-organise to form complex assembloids

Mature cell types in primary gastric organoids were scarcely detected, or absent, suggesting that organoid expansion alone is not sufficient to recapitulate full epithelial gastric function. In particular, spontaneously emerging parietal cells have never been reported, to date, in human primary organoids in expansion medium^15^, but only seen transiently in hPSCs^10,14^. Literature reports an overall difficulty in their identification in primary murine models as well^12^, but there are examples of murine-derived fundic organoids presenting parietal cells^13^.

Therefore, we designed a system to allow the self-aggregation of human gastric organoids in more complex multi-regional structures, defined as assembloids, where cells can reach maturity and functionality (**Fig.2a**). To facilitate organoid self-assembly, whilst maintaining the regional and orientational information, we designed a customized PDMS-based culture well (**Extended data Fig.2a**). Briefly, fully grown organoids were released from Matrigel, resuspended in collagen I hydrogel, and cultured in floating conditions, after assessment of the ideal gel concentration at 0.75 mg/mL (**Extended data Fig.2b, Supplementary Video 1**). We firstly defined the self-assembly conditions for each region (hereafter noted as single-region assembloid, SRA) and then multi-regional, in the order fundus-body-antrum (hereafter noted as multi-regional assembloid, MRA), to discern whether merely increasing complexity facilitates increased functionality, or whether multi-regionality is needed to enhance this further. The related organoid/collagen structures were cultured in suspension for 10 days, allowing contraction and self-aggregation (**Extended data Fig.3a-b**). By day 4, syncytia between organoids could be observed (**Fig.2b, Extended data Fig.3c-d**). By day 10, the self-assembled MRA and SRA structures were fully formed and used for subsequent analysis (**Fig.2c,f**). SRA transversal section showed a layer of cells polarized towards a unified epithelial lumen (**Fig.2d**). The expression of MUC5AC, MUC6, and CHGA confirmed the SRA gastric identity (**Extended data Fig.3e**). Whole mount immunostaining analyses on SRA showed the presence of a continuous shared lumen, with luminal apertures connecting to protrusions from the main assembloid body (**Fig.2e, Extended data Fig.3f**). These epithelial gastric gland-like structures were characterised by apicobasal cell-polarity determined by luminal F-ACT and basal b4-INT (**Fig.2e**). Remarkably, immunofluorescence analysis on the MRA demonstrated the preservation of regional identity within the assembloid system. PDX1 was found on the antral side of the MRA, together with chromogranin A and somatostatin (SST) positive cells. Additionally, the MRA revealed high expression of MUC5AC, indicating the presence of active mucin-producing cells. Enterochromaffin-like cells were identified throughout the MRA by the presence of CHGA, while PGC positive cells were exclusively identified on the fundic side (**Fig.2g, Supplementary Video 2-3**) (Double channel image of Nuclei and PGC only in **Extended data Fig.3g**).

**Fig.2.**
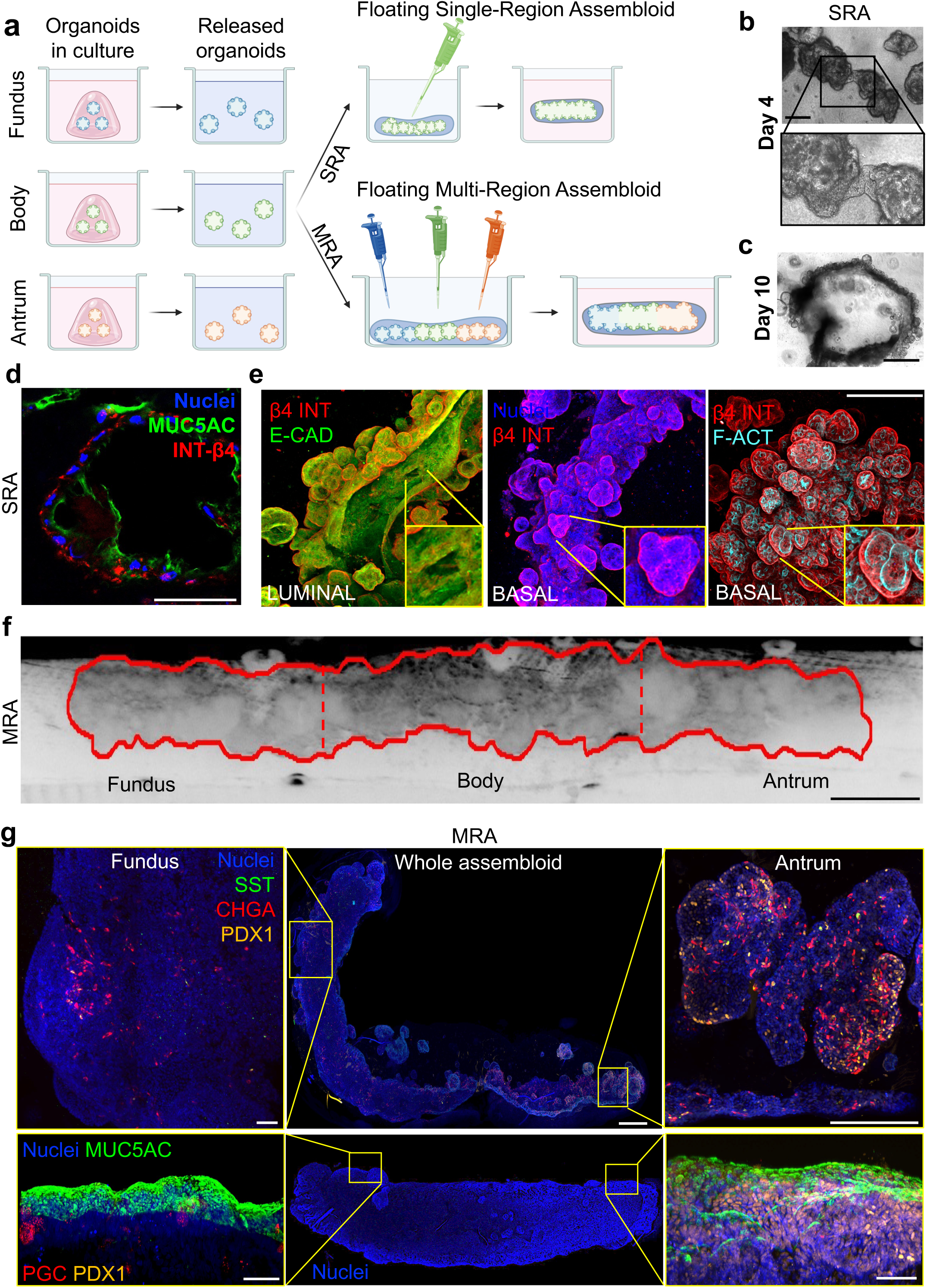
Gastric assembloids generation and characterization. **a**, Schematic representation of Single-Region Assembloid (SRA) or Multi-Regional Assembloid (MRA) generation. Created with BioRender.com. **b**, Bright field image of syncytia formation at day 4 of floating collagen I hydrogel culture. Scale bar 200 μm. **c**, Bright field image of 10days SRA. Scale bar 1 mm. **d**, Immunofluorescence of sectioned gastric SRA showing mucin 5AC (MUC5AC) in green, Integrin-b4 (INT-b4) in red, and nuclei in blue (Hoechst). Scale bar 50 μm. **e**, Whole mount images of gastric SRA showing Integrin-b4 (INT-b4) in red, and E-cadherin (E-CAD) in green in the upper panel (Luminal); nuclei in blue (Hoechst) and Integrin-b4 (INT-b4) in red in the middle panel; Integrin-b4 (INT-b4) in red and F-actin (F-ACT) in cyan in the lower panel (External). Images processed with IMARIS. Scale bar 500 μm. **f**, Bright field image of 10 days MRA. Scale bar 1 mm. **g**, Whole mount immunofluorescence of gastric MRA showing nuclei in blue (Hoechst), Somatostatin (SST) in green, Chromogranin A (CHGA) in red, Pancreatic and Duodenal homeobox 1 (PDX1) in yellow; or nuclei in blue (Hoechst), Mucin 5AC (MUC5AC) in green, Pepsinogen C (PGC) in red, Pancreatic and Duodenal homeobox 1 (PDX1) in yellow. Central column showing an overview of the whole tube structure (scale bar 500 μm). Left and right columns showing high magnification views of Fundus and Antrum ends (scale bar 50 μm).

### Cross-talk pathways arise in multi-regional gastric assembloids

Following RNA sequencing analysis, PCA of fundus, body, and antrum from tissue, organoids, SRA and MRA showed that all the *in vitro* systems clustered separately from the native tissue samples (**Fig.3a**). Interestingly, antrum SRA clustered closely to organoid samples, while antrum MRA clustered closely to antrum tissue on PC2, pointing to a shift in expression of the assembloids becoming increasingly similar to the *in vivo* condition only after co-culture. A separate PCA analysis on SRA and MRA only, showed how these two groups cluster separately (**Extended data Fig.4a**). Overall, the MRA displayed a variety of differentially expressed genes (DEGs), like the epithelial maturation marker *PHGR1*^25^, when compared to either organoids (**Extended data Fig.4b**) or SRA (**Extended data Fig.4c**), suggesting an advantage in the multi-regional complex system^26^. Looking at selected gastric targets, *PDX1* was confirmed antral specific in all sample types. Stem cell marker *LGR5* showed a reduction in MRAs compared to other *in vitro* systems. Endocrine cell markers (*Gastrin*, *Somatostatin* and *Ghrelin*) were found to be present in the assembloids coherently to their regional expression in the tissue. Parietal cell markers (*ATP4A* and *ATP4B*) mRNAs were lowly expressed in all in *vitro* systems when compared with gastric tissue (**Fig.3b**), although the RNA expression is still present (**Zoom Fig.3b**). Of note, a daily RT-qPCR during the 10-day MRA formation showed how the *ATP4B* mRNA spiked at around day 6, then decreased by day 8 (**Extended data Fig.4d**). Therefore, this temporal dynamic is coherent with the low detection at experimental endpoint.

**Fig.3.**
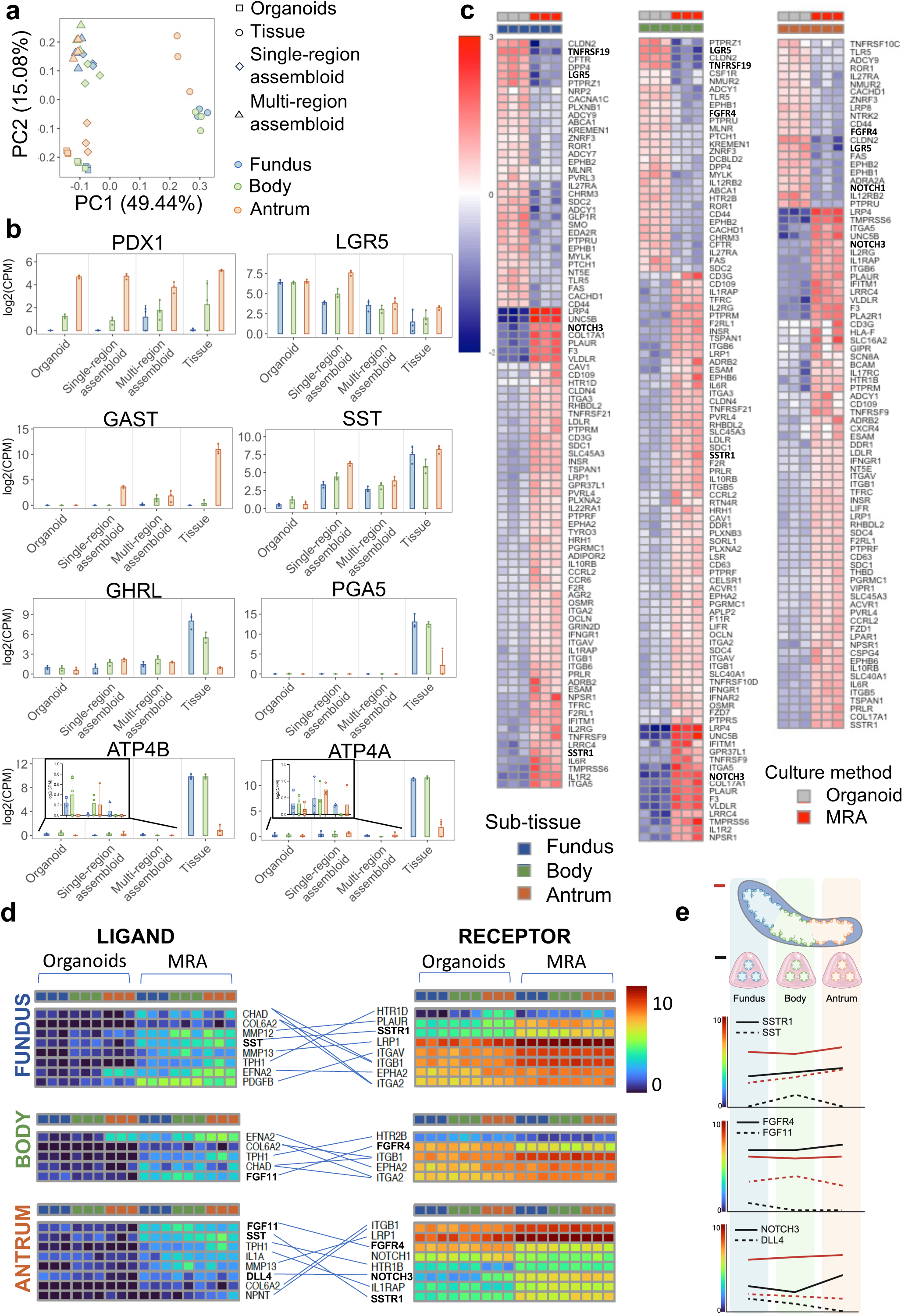
Gastric assembloid bulk RNA sequencing analysis. **a**, Principal component analysis (PCA) of RNA-sequencing samples from Fundus (blue), Body (green), and Antrum (red) organoids (squares), tissue (circles), SRA (rhomboid) and MRA (triangle). (n=3 for biological replicates). **b**, Expression of typical gastric markers in organoids, SRA, MRA and tissue in Fundus (blue), Body (green), and Antrum (red). Circles indicate single data points, mean ± SD (n = 3 for biological replicates) CPM: counts per million. **c**, Hierarchical clustering of receptors that are differentially expressed genes (DEGs) between organoids and MRA in the three regions. Colorbar indicates centered log2(CPM+1). **d**, Ligand-receptor couples expression in Fundus, Body, and Antrum in organoids vs MRA. The receptors are DEGs in the indicated subtissue between organoids and MRA. Corresponding ligands are DEGs between indicated subtissue between organoids and MRA, plus they are not expressed in the organoids of one subtissue (specified on the left). Colorbar indicates non-centered log2(CPM+1). **e**, Graphics summarizing the most relevant ligand-receptor couples emerging from sequencing data analysis in fundus, Body, and Antrum within the MRA (red line) and in the organoids (black line). Solid lines for receptors, dashed lines for ligands. Created with BioRender.com.

Subsequently we performed a ligand-receptor couples’ analysis to identify signalling pathways, arising in MRA when compared to simpler organoid systems, that might be responsible paracrine effects inducing cell differentiation. First, the differentially expressed receptors identified highlighted a marked difference expression between organoids and MRA of the different regions (**Fig.3c**). Of particular interest was the reduced expression of stem cell markers *LGR5* (in all the regions) and *TROY* (a.k.a. *TNFRSF19*, in body and fundus, where it is physiologically expressed) in the MRA compared to organoids, suggesting a loss of stemness in favour of a more differentiated phenotype, as previously reported in primary gastric organoids^15^. In addition, it has been reported that the Notch pathway regulates epithelial cell differentiation in the antral stomach through the receptors *Notch1* and *Notch2*, by promoting progenitor cell proliferation and inhibiting differentiation^27^. We found the presence of a reduced expression of *NOTCH1* in the antral region of the MRA compared to antral organoids. Of note, body and antrum regions in the MRA presented a decreased expression of *FGF4R*, which mediates cell proliferation in gastric cancer^28^. On the other hand, several protein tyrosine phosphatase receptors such as *PTPRS*, *PTPRF*, or *PTPRU*, responsible for cell differentiation regulation, were more highly expressed in the MRA compared to organoids. Finally, it is important to mention the increase in expression of *SSTR1* (*Somatostatin receptor*) in MRA. Unexpectedly, we faced the presence of inflammation-related markers.

We then performed a deeper analysis focusing upon ligand-receptor interactions, with the aim of highlighting the interactions that were absent in the organoid system and brought a significant change in receptor expression in MRA thanks to co-culture. In particular, we analysed the differential ligand expression between MRA and organoids in each gastric region, further shortlisting the ligands based upon where they were not expressed in the organoids (**Fig.3d**). Interestingly, many of the ligands identified were involved in extracellular matrix (ECM) remodelling. Functional enrichment analysis within Reactome database was performed to confirm this observation (**Extended data Fig.5**).

Finally, we focused on the most relevant pairs within the list. Fundic and antral organoids showed negligible expression of somatostatin, while its expression increased in MRA, particularly at the antral level. This was coupled with the increase in its receptor *SSTR1* at the MRA level, as previously mentioned. Furthermore, *FGF11* was found not to be expressed in body and antral organoids, but expressed in the MRA (**Fig.3e**). *NOTCH3* (receptor of *DLL4* ligand) increased in expression in the MRA compared to the single organoids. The analysis was repeated to investigate potential differences between SRA and MRA. The number of receptors identified as DEGs in fundus and body was approximately halved, while a higher number of DEGs was present in the antral comparison that brought to the identification of some more receptor-ligand interactions (**Extended data Fig.6a-b**). These data confirm that relevant ligand-receptor pairs arise when multi-regions are in communication in a single system, as in the MRA, allowing for cell cross-talk.

### Gastric assembloid system promotes parietal cell differentiation and function

Having established an enhanced morphological complexity with increased gastric epithelial differentiation and crosstalk, we then focused on assessing the gastric functionality of the MRA. We performed thiol organoid barcoding *in situ* mass cytometry (TOB*is* MC)^29,30^ to characterise the post-translational / signalling profile of MRA and organoids at single cell resolution; an optimized technique for high dimension panels^31,32^. We compared a panel of 30 proteins in the three MRA regions compared to the individual organoid cultures (**Extended data Fig.7**). To aid cell-type-specific analysis, fundic organoids were labelled with RFP, and body organoids were marked by GFP (**Extended data Fig.9a-b**). Antral organoids were not labelled due to the presence of markers that allow their unique identification (PDX1) (**Extended data Fig.9c**). PCA of protein signals clearly resolved organoid and MRA signalling (**Fig.4a**).

**Fig.4.**
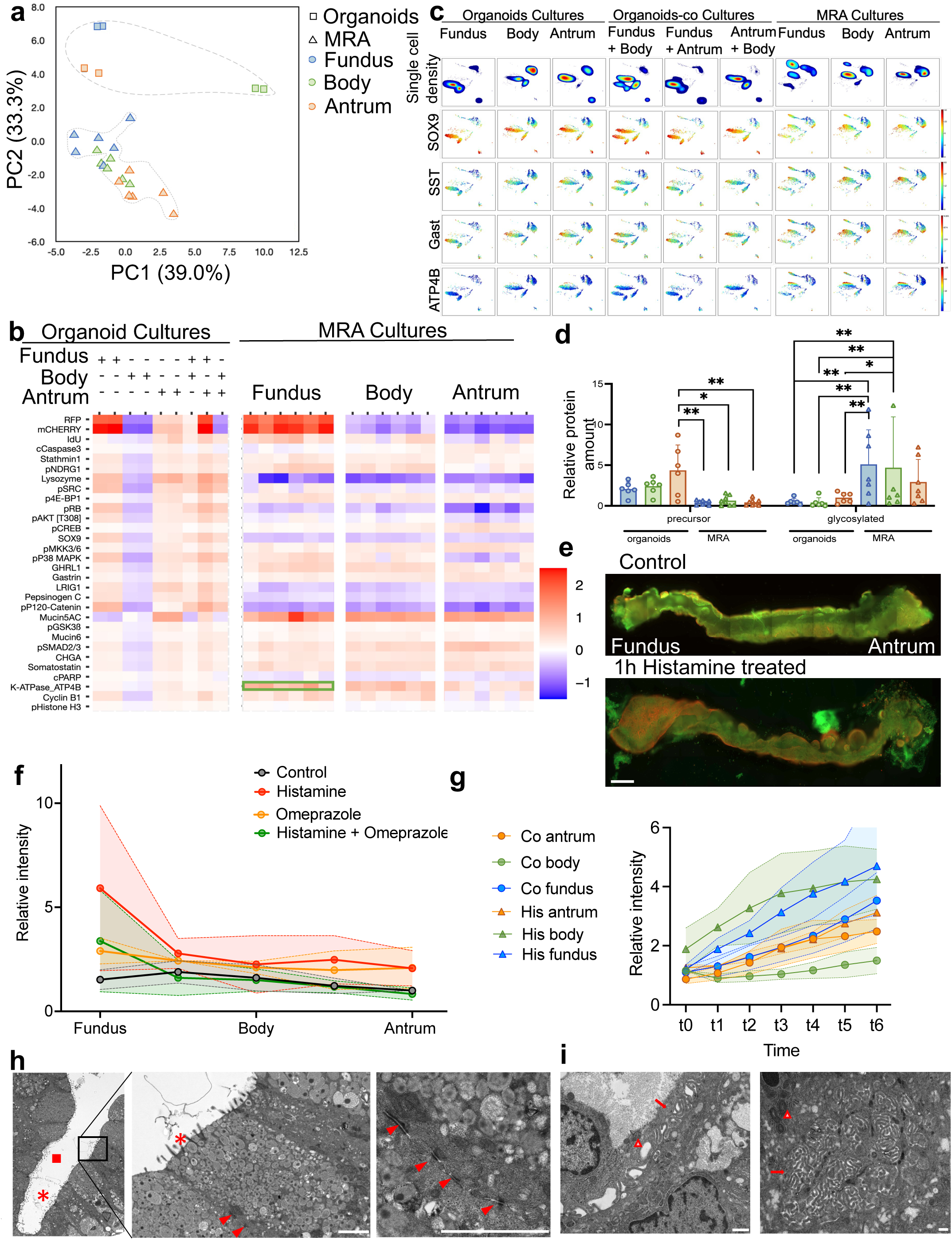
Mass cytometry and functional analysis of multi-regional assembloids. **a**, Principal component analysis (PCA) of TOB*is* MC data from Fundus (blue), Body (green), and Antrum (red) organoids (squares), and MRA (triangle). (n=2 for biological replicates for organoids, n=6 for biological replicates for MRA). **b**, Heatmap showing the abundancy of selected markers by TOB*is* MC analysis in organoids cultures compared to MRA. Green box highlighting the ATP4B expression in Fundus MRA. **c**, UMAPs of selected markers (SOX9, SST, GAST, ATP4B) from TOB*is* MC analysis in organoids cultures compared to MRA. **d**, ATP4B protein expression in Fundus, Body, and Antrum in organoids versus MRA using WES analysis for precursor (47-53kDa) and glycosylated protein (60-80kDa). Circles indicate organoids single data points, squares indicate MRA single data points, mean ± SD (n=6 biological replicates for organoids, n=7 biological replicates for MRA). Two-way Anova (adjusted p-value * < 0.05. ** < 0.01). **e**, Acridine orange staining of MRA with or without Histamine treatment (100 μM). Excitation wavelength 488 and 568 nm. Scale bar 500 μm. **f**, Acridine orange 568/488nm ratio quantification of Fundus, Body, and Antrum in MRA, normalized to Antrum-control. Control, Histamine treatment (100 μM), Omeprazole treatment 100 μM, Histamine (100 μM) + Omeprazole (100 μM). Circles indicate single data points, mean ± SD (n = at least 2 for technical replicates – 2 patients). **g**, Acridine orange 568/488nm ratio timelapse quantification of single cells from Fundus, Body, and Antrum in MRA. Control (circles), Histamine treatment (100 μM - triangles), Data points indicate average of 5 to 7 cells per each point. mean ± SD (1 patient). **h**, Transmission electron microscopy (TEM) images of MRA. Grey circles within cells indicating mucin-filled granules. Scale bar 2 μm. ▪ = gland, * = microvilli, ▴ = desmosomes. **i**, TEM images of MRA showing parietal cells. Scale bar 500 nm. △ = mitochondria, → = canaliculi.

Overall, the MRA demonstrated higher protein levels from differentiated cell types (**Fig.4b**)^33^. Stem cell markers (SOX9) were found to be less expressed in the assembloid (**Fig.4c**). Antral specific markers such as SST and GAST were mostly found in the antral side of the multi-regional assembloid. Interestingly, parietal protein ATP4B expression was significantly higher in MRA compared to organoids, with physiological expression on the fundus and body of the assembloid (**Fig.4c**). Of note, the mere coculture of organoids in Matrigel was not enough to induce a shift of expression from a organoids signature to a MRA one. Indeed, plotting the normalized reads corresponding to the different conditions, we demonstrated the significant difference between MRA versus organoids or organoids co-culture (**Extended data Fig.9**): stem markers LRIG1, stathmin, SOX9, were all downregulated in MRAs compared to other conditions, underlying the promoted cell differentiation in the assembloids. On the other hand, gastric differentiation markers GAST, SST, Ghrelin, MUC5AC were all upregulated in MRAs compared to the other conditions. Also, GAST was regionally overexpressed in the MRA antrum compared to fundus, ghrelin was more expressed in the MRA fundus/body compared to antrum, and mucin was equally expressed through the MRA regions, making the model reliable and the regionality preserved. Moreover, ATP4B protein was significantly overexpressed in MRA compared to both single organoids and organoid regional duplets (organoid co-cultures), and the expression was confirmed to be higher in the MRA fundus compared to the antrum (**Extended data Fig.9**). To investigate whether MRA are able to produce and secrete gastric acid, the glycosylated functional form of ATP4B (as found in parietal cell membranes) was evaluated by WES protein analysis. Strikingly, we verified a significant higher level of glycosylated ATP4B protein amount in the MRA fundus and body compared to its antral side, and each regional organoid, compared to the precursor form of the protein (**Fig.4d**). At the functional level, the presence of acid production mediated by the activity of the glycosylated functional form of ATP4B (as found in parietal cell membranes) in the complex of ATPase H+/K+ pump was evaluated by live staining with acridine orange (AO), a dye excited when the pH is ≤3.5. We observed a strong increase in fluorescence in MRA treated with histamine, which physiologically stimulates parietal cells to secrete acid with analysis performed at endpoint (1h histamine stimulation) (**Fig.4e**). We compared 4 conditions: i) control stained with AO, ii) treated 1h with histamine, iii) treated with omeprazole (chemical inhibitor of the pump), iv) treated with omeprazole and histamine. We observed no significant difference in the 3 regions of the MRA control (i), and no difference in the omeprazole-treated assembloid (iii). In contrast, the addition of histamine (ii) induced an increase of fluorescence intensity, indicating higher acid secretion. Notably, the combination of histamine and omeprazole (iv) showed a signal reduction compared to histamine treatment alone (ii) (**Fig.4f**). This analysis was performed considering the maximum fluorescence intensity reached in each region and compared among conditions. Given the larger structures in our experimental condition, we performed a timelapse analysis by area (fundus, body, antrum) post drug administration at intervals comparable to what previously published^10^ (**Extended data Fig.10a – Supplementary Video 4-5-6**).

Acridine orange accumulation was observed in a canalicular-type pattern in parietal cells, in a similar fashion to what already described^10^ (**Extended data Fig.10b**). Moreover, we examined single-cell progression through a 1h timelapse microscopy, comparing control condition to histamine stimulation (**Fig.4g**). Consistently, we detected an increase of the signal ratio, with higher signal in cells on the fundic side, showing the highest ratio upon histamine stimulation.

We then performed transmission electron microscopy (TEM) analysis on the 3 regions of the MRA to morphologically confirm the presence of the functional cell types. We identified defined gastric glands, with a mature epithelial monolayer of mucus-secreting cells. We confirmed the presence of microvilli on the apical domain, as well as desmosomes mediating cell adhesion on the lateral domains (**Fig.4h**). Importantly, we identified the presence of parietal cells in the MRA body and fundus regions, showing large mitochondria and canaliculi cytoplasmic network (**Fig.4i**). Taken together, these data suggest that the multi-regional assembloid system favours enhanced gastric parietal differentiation with the appearance of complex gastric specific functionality.

### Assembloids show gastric function preservation upon *in vivo* transplant

To further promote assembloid differentiation and prove its survival, we evaluated its response *in vivo.* To accomplish this, we subcutaneously implanted the SRA and MRA into immunodeficient NSG mice. Silicone O-rings were used to spatially track the implanted assembloids (**Fig.5a**). The implants were retrieved 4 weeks post transplantation where all transplanted segments were viable and surrounded by host mesenchyme and vasculature (**Fig.5b**). Furthermore, transplanted assembloids showed larger structural features compared to *in vitro* assembloids, including a lumen with a significant increased volume and the persistence of gland-like regions. For a comprehensive morphological evaluation, explants were characterized by microfocus computed tomography (Micro-CT) and whole-mount immunofluorescence. Three-dimensional reconstruction of the SRA Micro-CT demonstrated preservation of a shared large lumen and connected gland-like structures opening into the lumen (**Fig.5c, Supplementary Video 7-8**). The annotated seconds in Figure 5C refer to the corresponding frames in the original video showing the 3D reconstruction of the structure (**Supplementary Video 8**). Immunofluorescence confirmed the persistence of a polarised epithelium with basal lamina (β4-integrin), and secretion of MUC5AC in the lumen (**Fig.5d**), suggesting maintenance of gastric epithelial cells identity.

**Fig.5.**
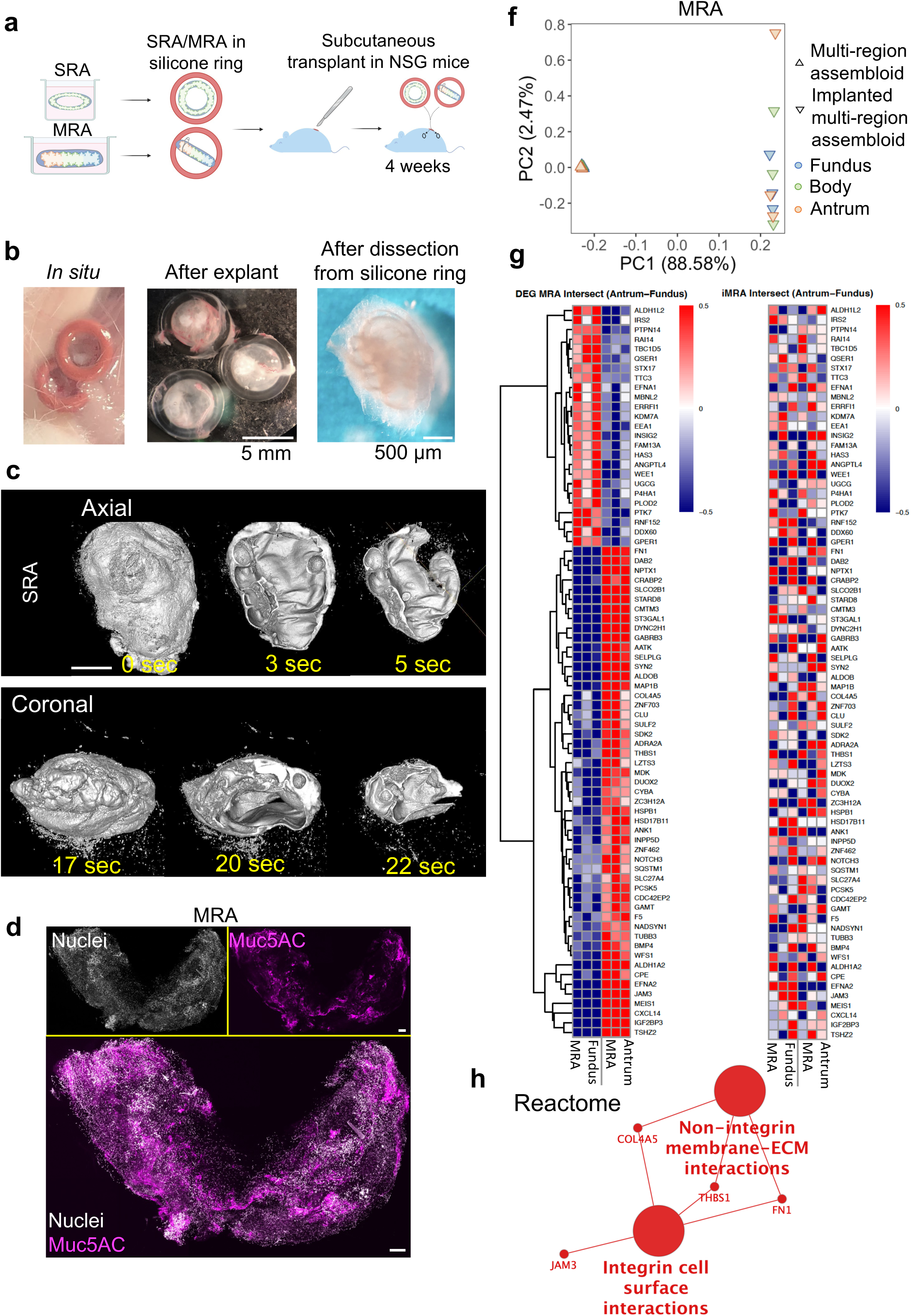
In vivo implantation of single- and multi-regional gastric assembloids. **a**, Schematic representation of assembloid implantation workflow. Created with BioRender.com. **b**, Post-implantation assembloid harvesting: sub-cutaneous location (left), after explanation (central, scale bar 5mm), isolated assembloid (right, scale bar 500 μm) (n=3 for biological replicates). **c**, Microfocus computed tomography (Micro-CT) of transplanted SRA from axial and coronal view. Scale bar 500 μm. **d**, Whole mount images of gastric SRA showing nuclei in blue (Hoechst), Mucin 5AC (MUC5AC) in red, Integrin-b4 (INT-b4) in green. Scale bar 500 μm. **e**, Principal component analysis (PCA) of RNA-sequencing samples from Fundus (blue), Body (green), and Antrum (red) in MRA (△) and post-implantation MRA (iMRA, ▽). (n=3 for biological replicates). **f**, Hierarchical clustering of DEGs (FDR<0.05, fold change > 1.5) between Antrum and Fundus in MRA (left), and expression of the corresponding genes in iMRA (right). (n=3 for biological replicates). Color bar refers to centered log2(CPM+1). **g**, Functional enrichment analysis of genes displayed in (F) within Reactome database. Larger dots indicate enriched functional categories, small dots genes belonging to the linked categories.

RNA sequencing was performed on post-implantation MRA (iMRA). PCA between MRA and iMRA showed a distinctive separation between the two groups of samples (**Fig.5e**). Further analysis comparing antral and fundic expression within the assembloids showed significant heterogeneity in the iMRA, ascribable to the biological variability in an *in vivo* environment. (**Fig.5f**). Overall, these heatmaps illustrate the DEGs between Antrum and Fundus that are DEG both in MRA and in iMRA, with gene order established by hierarchical clustering. Notably, the DEGs analysis comparing antrum vs fundus in MRA and iMRA showed a decreased stem cell marker expression in fundus iMRA compared to antrum, suggesting a higher fundic-phenotype differentiation *in vivo*. Additionally, further extrapolated DEGs among a curated list of gastric-specific genes of antrum vs fundus in MRA and iMRA showed a decreased stem marker expression in fundus iMRA compared to antrum, suggesting a higher fundic-phenotype differentiation *in vivo*, as well as functional markers such as aquaporin 5, physiologically upregulated in the antrum side of the iMRA (**Extended data Fig.11**). Reactome functional enrichment analysis revealed a general increase in ECM-interacting proteins (**Fig.5g**).

### Multi-regional gastric assembloids model HNF4a/PMM2 genetic disease

Endoscopic biopsies from two patients with PMM2-HIPKD-IBD were used to derive gastric organoids and implement our MRA system, hereafter ΔMRA (Δ*PMM2* MRA).

Fundus, body and antrum organoids were successfully derived from endoscopic biopsies of both patients (**Fig.6a**). For one of the two patients, a biopsy was collected directly from a hyperplastic polyp itself, allowing the derivation of organoids from this fourth anatomic location (**Extended data Fig.12a-b**). Upon initial morphological comparison with organoids derived from healthy control (HC), both organoid conditions shared a cystic and round appearance. However, Δ*PMM2* organoids exhibited higher abundance of bud-like protrusions at similar passage (**Fig.6a**), and were significantly larger compared to the HC counterpart (**Fig.6b**). Although the number of KI67+ proliferative cells was comparable among all organoid lines (**Fig.6c**), in the ΔMRA we identified focal regions with significantly enhanced proliferation, particularly within the antral side (**Fig.6d and Extended data Fig.12c**). This phenotype was even more pronounced when analysing polyp-derived organoids, where whole PDX1+ protruding buds were highly positive for K67 (**Fig.6e**). H&E staining of ΔMRA slices revealed the presence of cell aggregates exclusively on one side of the MRA (**Fig.6f**). These structures, representing hyperplastic protrusions, mirror the foveolar hyperplasia seen in patient biopsy samples, as highlighted by histopathological examination with toluidine blue (**Fig.6g**). TEM analysis of the corresponding tissue biopsies revealed a thinner layer of mucin granules in mucous-producing cells compared to normal conditions. Moreover, mucin granules in the hyperplastic regions were less electron-dense, synonym of a less mature state (**Fig.6g**). We found a corresponding reduction in the amount of MUC5AC Δ*PMM2* organoids (**Fig.6h**), and in the hyperplastic regions of ΔMRA (**Fig.6e**). RT-qPCR indicated a significant increase in the gene expression of *MUC5AC* in MRAs compared to organoids in HC, aligned with an increase in cell differentiation, but this was absent ΔMRAs (**Fig.6i**).

**Fig.6.**
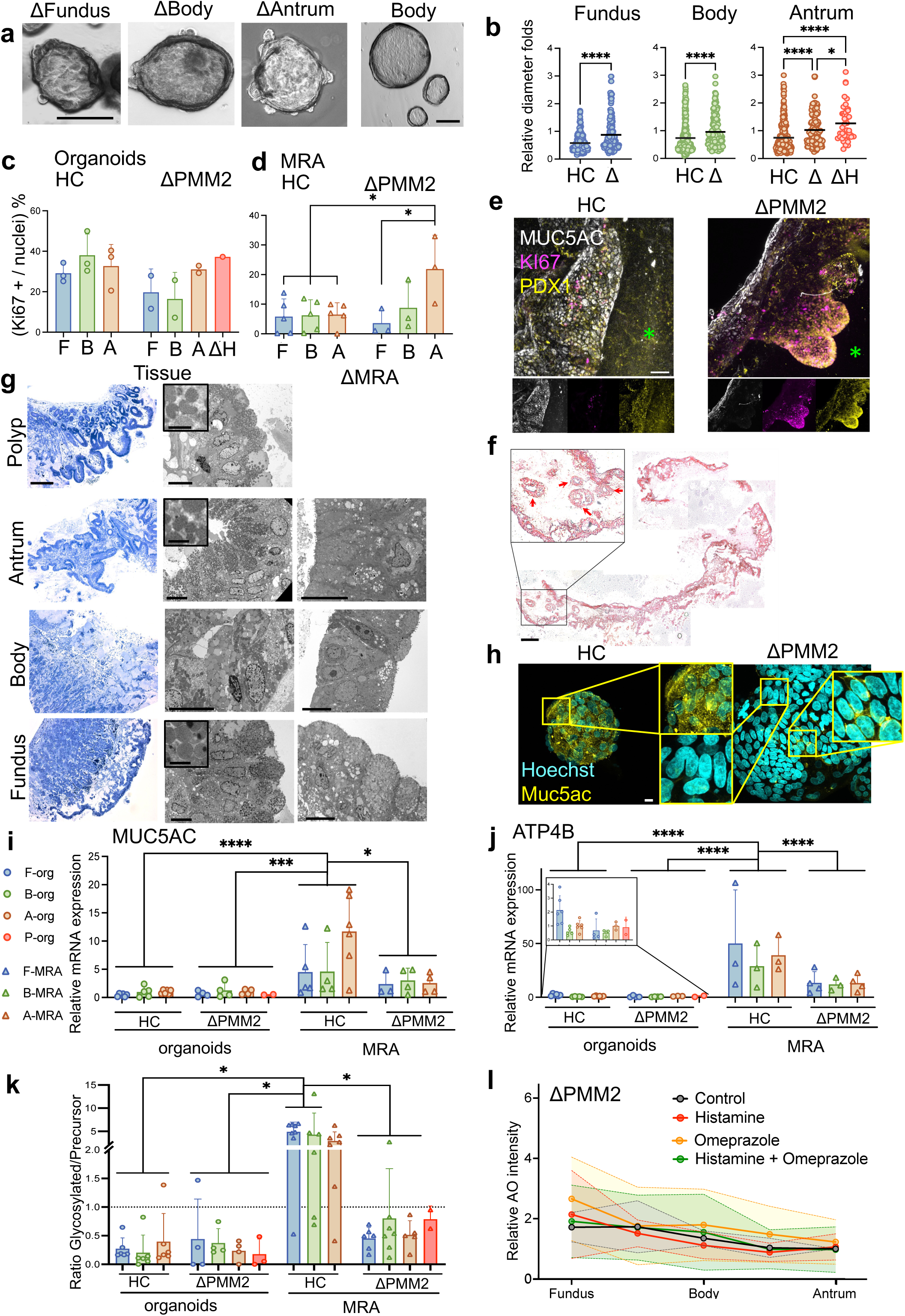
Disease modelling of HNF4a/PMM2 genetic condition with multi-regional gastric assembloids. **a**, Bright field image of representative organoid line for Fundus, Body, Antrum of ΔPMM2 showing the formation of spherical organoids with visible buddings within 7 days starting from single cells. Comparison to 7 days HC Body, Scale bar 100 μm. **b**, Evaluation of organoid diameter of fully grown organoids at day 7 of culture (3 organoid lines derived from patients not affected by the mutation (HC) and 2 organoid lines derived from patients affected by the mutation (ΔPMM2, only one for Hyperplastic protrusion condition). Values are displayed as relative fold to the average of the Antrum-mutant. Mean shown as a bar, dots represent single organoids measurement. HC = Healthy control, Δ = ΔPMM2, ΔH = ΔPMM2 hyperplastic protrusion). T-test analysis to analyse differences for each region (HC fundus vs Δ fundus; HC body vs Δ body). One-way Anova to compare HC Antrum vs ΔAntrum vs ΔH. Adjusted p-value * < 0.05, **** < 0.0001. **c**, Ki67 positive cell count over total nuclei percentage in whole mount staining of 3 HC organoid lines and 2 ΔPMM2 organoid lines. Mean ± SD. One dot for each patient. F = fundus, B = body, A = antrum, ΔH = Hyperplastic protrusion. One-way Anova – non significant **d** Ki67 positive cell count over total nuclei percentage in whole mount staining of MRA regions. 3 HC patients (n=5 replicates, left side of the graph) and 2 ΔPMM2 (n=3 replicates, right side of the graph). F = fundus, B = body, A = antrum. Adjusted p-value from the one-way Anova * < 0.05. **e**, Whole mount immunostaining of antral side of MRA and ΔMRA made of ΔH organoids: KI67 in magenta, PDX1 in yellow, MUC5AC in white. (scale bar 20 μm). **f**, Reconstruction of H&E-stained section of ΔMRA. Arrows indicating cell aggregates (Scale bar 500 μm). **g**, Toluidine blue staining (scale bar 200 μm) and matching TEM images of ΔPMM2 tissue samples (Scale bar 10 μm). Zoom areas displaying mucin granules (Scale bar 2 μm). (n=1). **h**, HC patient and ΔPMM2 patients derived organoids whole mount staining. Nuclei in cyan, MUC5AC in yellow. Green asterisks indicate the lumen. Scale bar 50 μm. **i**, RT-qPCR for Muc5ac, Organoids vs MRA, HC vs ΔPMM2. (HC organoids 5 replicates, 3 patients; ΔPMM2 organoids 4 replicates, 2 patients – ΔH present only in duplicate for 1 patient; HC MRA 5 replicates, 3 patients; ΔMRA 4 replicates, 2 patients. Adjusted p-value from the two-way Anova (* < 0.05, *** < 0.001, **** < 0.0001). **j**, RT-qPCR for Atp4b, Organoids vs MRA, HC vs ΔPMM2 (HC organoids 6 replicates, 3 patients; ΔPMM2 organoids 4 replicates, 2 patients – ΔH present only in duplicate for 1 patient; HC MRA 3 replicates, 3 patients; ΔMRA 4 replicates, 2 patients. Adjusted p-value from the two-way Anova (**** < 0.0001). **k**, Glycosylated over precursor ATP4B protein expression ratio in Fundus, Body, and Antrum in organoids versus HC MRA versus ΔMRA using WES analysis. Circles indicate organoids single data points, triangles indicate MRA single data points, mean ± SD (n=6 biological replicates for HC organoids, n=4 biological replicates for ΔPMM2 organoids, n=7 biological replicates for HC, n=6 biological replicates for ΔMRA). Adjusted p-value from the two-way Anova (* < 0.05, **<0.01). **l**, Acridine orange 568/488nm ratio quantification of Fundus, Body, and Antrum in ΔMRA, normalized to Antrum-control. Control, Histamine treatment (100 μM), Omeprazole treatment 100 μM, Histamine (100 μM) + Omeprazole (100 μM). Circles indicate average of single patient data points, mean ± SD (n = 3 for technical replicates – 2 patients).

Subsequently, we focused on ATP4B, given that its functional form relies on its N-glycosylation. The expression increase of *ATP4B* was increased in MRAs compared to organoids. This increase was less in Δ*PMM2* MRAs (**Fig.6j**). The ATP4B subunit is translated in a first peptide of 34kDa, and undergoes a process of post-translational modification to generate the functional protein that is translocated to the membrane for acid production^34,35^. The process produces an intermediate precursor of 43-57kDa, which is glycosylated generating a span of proteins of different molecular weights ranging from 60 to 80kDa, allowing protein transportation and retention to the membrane. We were able to identify the three protein forms in both HC and ΔPMM2 patients in three separate peaks at the indicated molecular weights (**Extended data Fig.12e**), and we calculated the ratio of quantified values of glycosylated to precursor protein as an index of N-glycosylation activity. As previously mentioned, in the HC patients we detected enhanced glycosylation activity in MRA compared to organoids. This was significantly decreased in ΔMRA, strongly indicating a reduced N-glycosylation in the context of the PMM2 variant (**Figure 6k)**.

Lastly, histamine stimulation of ΔMRA did not result in a pH change in acridine orange assay, confirming the reduction in functional glycosylated ATP4B (**Fig.6l**). Overall, these data suggest that the MRA system can model the gastric epithelial dysfunctions observed in PMM2 patients.

## Discussion

This study focuses on the establishment of region-specific gastric organoids and their engineering into a multi-regional assembloid system to model the human gastric epithelium.

The stomach is a complex organ with distinct anatomical regions defined antrum, body and fundus^26,36^. PDX1 has been identified as a marker of developing gastric antrum in mice^21^, as well as being expressed in iPSC-derived gastric organoids^22^. Recognising the important patterning role of PDX1 during gastric regionalisation^37^, we verified its presence at fetal stage PCW10, and its persistence in gastric tissue derived from infants and children. We showed conservation of PDX1 expression in the antral lines we characterised, confirming the ability of primary derived organoids to maintain the spatial information of the gastric region of origin.

Our results demonstrate how region specificity is maintained in the organoid lines. These findings were confirmed by bulk RNA sequencing and protein validation of the organoids, and were independent from any extrinsic signalling, since organoids from different regions were cultured in the same expansion medium. This simpler model is, however, still lacking mature gastric functions such as acid secretion.

Assembloids provide a novel strategy to recapitulate complex tissue interactions *in vitro*. Multiple organoids can be combined to form more complex models, enabling the study of multiple cell types simultaneously^38^. When we increased the complexity of our system, by building single-region assembloids, we observed a degree of cell maturation demonstrating a clear advantage over the organoids. However, this SRA model lacks cross-communication between the different stomach regions. The degree of differentiation induced in SRA was further enhanced in multi-regional assembloids, due to the biochemical cross-interaction between the regions. We were able to assess the regional marker preservation in each region of our MRA, as well as gastric identity retained with the production of mucins and gastric-specific proteins (i.e., somatostatin, gastrin). Our system allows the interaction of the 3 regions with the establishment of a shared lumen, defined apicobasal cell polarization, and gastric gland-like protrusions.

Notably, PDX1 expression remained confined to the antrum region, indicating the absence of organoid migration along the assembloid, during the organoid self-assembly. This suggests that gastric organoids have the capacity to retain the spatial information and identity of the region of origin also in a more complex system. Transcriptomic analyses revealed that the native tissue samples, organoids, and assembloids cluster separately, indicating differences between *in vitro* systems and the native tissue. However, MRA demonstrated a closer transcriptomic profile to the tissue cluster, compared to organoids alone. Our system showed overall increased level of differentiation markers, confirmed at transcriptomic and proteomic level. Interestingly, MRA shows expression of mature mucin-secreting and endocrine cells, whilst expressing a stem-progenitor compartment at a comparable level to the tissue of origin. This is in contrast to organoids which retain a less differentiated transcriptomic profile. Also, the antrum transcriptomic profile appears to differ the most, suggesting that it is favoured in the multi-regional approach.

Ligand-receptor analysis highlighted differential expression patterns in MRA compared to organoids, providing insights into potential paracrine signalling pathways involved in differentiation. Coherently with the increase of differentiation markers in MRA, we observed a reduction in the expression of receptors that mediate a stem cell state (most notably LGR5). Of particular interest, we observed the increase in expression of the ligand somatostatin, particularly expressed in the antral region of MRA, with its receptor SSTR1, expressed over all the regions in the MRA. Somatostatin is a peptide hormone, produced by the Delta cells, which plays a significant role in controlling the function of parietal cells, thus regulating pH level, and maintaining the balance of digestive processes^39^. In addition, a number of inflammation-related receptors were identified. Importantly, these assembloid structures have a considerably larger scale than single organoids. Consequently, there might be a reduced medium diffusion in the inner part of the assembloid, or an accumulation of dead cells in the enclosed lumen, impeding their elimination.

Generating parietal cells in adult stem cell-derived gastric organoids has been challenging to date. We hypothesized that our MRA system could facilitate the emergence of this cell type. To identify the elusive parietal cell type, we adopted a single-cell proteomic approach.^30–32,40^ Cytometry by time of flight confirmed some of the transcriptomic findings in the MRA. In particular, at the proteomic level, stemness markers were reduced, alongside a reduction in proliferating stem cells as indicated by the MRA cell state profile. Concurrently, antral differentiation proteins such as gastrin and somatostatin increased in the antral region of MRA. The main finding of the proteomic analysis was the presence in the MRA fundic region of the gastric specific proton pump ATP4B, responsible for acid production. The presence of the active glycosylated form of the ATPase H+/K+ transporting subunit beta, as found in the parietal cells *in vivo*, was confirmed by WES analysis. Overall, these results provide very interesting insights. Firstly, the multi-regional assembloid does promote higher gastric differentiation compared to the single-region organoids. Secondly, the cross-communication between the regions is needed to promote this differentiation, as the duplets of single organoids that are cultured in the same Matrigel droplets but are not fused together, do not promote the same level of differentiation as the multi-regional assembloids.

We were able to assess acid secretion in the MRA by live imaging with pH-reactive dyes. Functional assessment using live imaging confirmed the presence of active proton pumps and acid production in the fundic and body regions of the assembloid. Strikingly, MRA responded to administration of histamine, a nitrogenous compound that stimulates gastric acid secretion, and omeprazole, a drug that inhibits H+/K+ ATPase proton pumps, reducing gastric acid secretion.

Moreover, we demonstrated *in vivo* survival of the assembloid system, upon transplantation in immunocompromised mice. Both SRA and MRA preserved the phenotypic characteristics of the assembloids with a larger lumen, identified by iodinated microfocus CT, and a complex epithelial wall with gastric mucous-secreting glands. The epithelium was able to maintain apicobasal polarity, and a regionalised gastric identity. Finally, using our MRA system we were able to produce a reliable patient-specific disease model. We provide the first clear evidence of defective N-glycosylation in the context of PMM2-HIPKD-IBD-associated *PMM2* variants in patient-derived samples with a context- and cell type-dependent effect. We provide evidence of impact on epithelial secretory function, which may be mechanistically linked both to the development of the unusual gastric pathological manifestations, and the inflammatory intestinal pathology more broadly. The *in vitro* framework we have develop to model the GI-specific effects of this disease, offers a platform for preliminary testing of therapeutic agents targeting this specific pathway, for example Epalrestat, which has been shown to activate PMM2 activity and is under trial for the management of PMM2-CDG^41^.

Overall, our work shows the successful isolation of gastric organoids derived from different regions of the stomach and their self-organisation capacity into complex multi-regional structure. These analyses demonstrate the occurrence of important gastric transcriptional changes upon self-assembly of the multi-organoid constructs. In particular, the floating culture system favours functional maturation of differentiated cell types. We believe that our gastric multi-regional assembloids can mimic complex *in vivo* stomach physiology, providing valuable insights for human gastric pathology and therapy investigation.

## Methods

### Ethics and licences

Human fetal stomachs were dissected from tissue obtained immediately after termination of pregnancy, in compliance with the bioethics legislation in the UK. Fetal samples were obtained from the Joint MRC/Wellcome Trust Human Developmental Biology Resource (HDBR) under Research Tissue Bank ethics approval UCL site REC reference: 18/LO/0822 - IRAS project ID: 244325. Samples were used for tissue characterisation.

All paediatric gastric tissue was donated by the patient, or person with parental responsibility, after informed consent was obtained by an independent research coordinator at the relevant study site, under the approved research license. Human gastric biopsies were collected and processed in compliance with the Human Tissue Act under licence 18DS02 by the NHS Health Research Authority, East of England, Cambridge Central Research Ethics Committee. Paediatric samples were sourced from endoscopic mucosal biopsies at Great Ormond Street Hospital NHS Foundation Trust. Samples were used for deriving organoid lines or for native tissue characterisation.

All animal care and procedures were performed in accordance with National Centre for the Replacement Refinement and Reduction of Animals in Research (NC3Rs) guidelines. Experimental work was undertaken under ethics approval governed by UK Home Office Project Licence PPL PDD3A088A and Personal Licence PIL I73E168C9.

### Gastric glands isolation

Gastric tissues were collected in ice-cold Advanced DMEM/F12 medium (Thermo Fisher Scientific, 12634) supplemented with 10 mM HEPES (Thermo Fisher Scientific, 15630080), 2 mM Glutamax (Thermo Fisher Scientific, 35050061), and 1% penicillin-streptomycin (Thermo Fisher Scientific, 15140122), defined as ADMEM+++, and processed immediately following collection. Complete set of biopsies was isolated from each patient (endoscopically or by sample resection after gastrectomy), comprising fundus, body and antrum samples, which were processed separately. Glands were isolated using an established chelation-mechanical dissociation protocol ^8,23^. Endoscopic gastric biopsies of clean mucosal layer (3-5mm diameter) were processed without further dissection. Tissue was washed in a petri dish with cold HBSS to remove debris and a glass coverslip was then used to scrape away the luminal mucus layer. Tissue was then cut into smaller pieces (1 mm diameter) with a scalpel, transferred to a 15 mL tube containing 10 mL of fresh HBSS and then washed vigorously using a 10 mL pipette pre-coated with 1% bovine serum albumin (BSA; A9418, Sigma-Aldrich) in Dulbecco’s phosphate buffered saline without calcium and magnesium ions (DPBS; D8537, Sigma-Aldrich). After washing, the pieces were allowed to settle, and the supernatant discarded. This process was repeated until the supernatant was clear. The mucosal pieces were then incubated in a chelating buffer made up of 5.6 mmol/L Na_2_HPO_4_, 8.0 mmol/L KH_2_PO_4_, 96.2 mmol/L NaCl, 1.6 mmol/L KCl, 43.4 mmol/L sucrose, 54.9 mmol/L D-sorbitol, 0.5 mmol/L DL-dithiothreitol, and 2mM ethylenediaminetetraacetic acid (EDTA) (all from Sigma-Aldrich), in Milli-Q water (18.2 mW/cm; Merck Millipore) for 30 minutes at 37°C on a planar shaking platform. EDTA was discarded and mucosal pieces washed in ice-cold DPBS with calcium and magnesium ions (DPBS++; Sigma-Aldrich, D8662). Washed mucosal pieces were then transferred to a new 10 cm petri dish on ice and hand pressure was applied to the lid in contact with the mucosal surface with a 3.5 cm petri dish to release the gastric glands. The released glands were collected from the dish with ice-cold ADMEM+++ and filtered through a 40 μm cell strainer and centrifuged at 200 g for 5 minutes at 4°C. Supernatant was aspirated and glands were resuspended in ice-cold undiluted Matrigel® Growth Factor Reduced (MGFR; Corning, 354230). Droplets of 30 μL were aliquoted into pre-warmed multi-well plates and incubated inverted for 20 minutes at 37°C to induce gelation. 10 μM Rho kinase inhibitor Y-27632 (Tocris, 1254) was added to the medium for the first 3-4 days. Gastric organoid medium was added and changed every 3 days. (**Supplementary Table 2**).

### Passaging gastric organoids

For expansion, organoids were passaged by enzymatic single cell dissociation. Organoids were retrieved from Matrigel with ADMEM+++ wash while working on ice. After retrieval from the well and centrifugation at 300 g for 5 min at 4°C, the organoid pellet was resuspended in 1 mL of TrypLE Express (Thermo Fisher Scientific, 12605010), incubated at 37°C for 5 minutes, and manually disaggregated with a P1000. TrypLE was quenched with 10mL ice-cold ADMEM+++ and centrifuged at 300 g for 5 minutes at 4°C. After aspirating the supernatant, single cells were then resuspended in MGRF at the desired split ratio (1:3 to 1:15) and plated in 30 μL droplets, in pre-warmed multi-well tissue culture plates. Rho kinase inhibitor Y-27632 was added to the medium for the first 24h.

### Generation of reporter cell lines for single cell experiments

Fundus-RFP (Red Fluorescence Protein) and body-GFP (Green Fluorescence Protein) reporter organoid lines were generated by electroporation based upon previously described methods (https://doi.org/10.1038/nprot.2015.088). Briefly, 10 μM Y-27632 was added to the culture medium 2 days prior and 1.25% DMSO was added 1 day prior to electroporation. Organoids were dissociated to single cells with Accumax (Thermo Fisher, 00-4666-56), filtered, centrifuged, and resuspended in 1 mL Opti-MEM (Thermo Fisher, 31985062) with Y-27632. PiggyBac vectors 1 μg/mL and piggyBac transposase 0.5 μg/mL were added to the cell suspension and transferred to a 2 mm electroporation cuvette and electroporated using a NEPA21 electroporator. Electroporated cells were transferred to a new 1.5 mL tube and 400 mL Opti-MEM with 10 μM Y-27632, centrifuged, supernatant aspirated, pellet resuspended in MGFR, and plated. After MGFR polymerisation, gastric medium with Y-27632 and 1.25% DMSO was added. DMSO was removed from the medium after 24 hours. Organoids with successful integration of the respective plasmids were selected with puromycin at day 5 after electroporation.

### Design and fabrication of custom culture well

A narrow linear culture well was designed to constrain the self-aggregation of organoids into a linear tube with fundus GOs at one end and antrum GOs at the other end. SolidWorks 2020 (Dassault Systems) was used to 3D design the well rectangular shape. The design was converted into coordinates using ideaMaker (Raise3D) to allow printing by an E2 desktop 3D printer (Raise3D). The mould was printed in polylactic acid (PLA) using fused filament extrusion deposition. Layer height was set to 0.05 mm to optimise smooth external surface of the walls of the mould. To complete the top of the pillars, an ironing function was applied, and the resulting structure was sanded by hand to smooth the surface. The resulting mould was used as a negative to cast a 10-well plate in polydimethylsiloxane (PDMS). A premixed 10:1 mixture of PDMS pre-polymer and curing agent solutions (Sylgard 184 kit, Dow Corning) was cast and cured at 40 °C overnight.

### Production of collagen I hydrogel

The following methodology was adapted from a previous published protocol^42^. Rat tail collagen I was used to prepare a collagen I hydrogel of physiological pH and salinity using the recipe in **Supplementary Table 3**. Collagen I concentrations of 0.75 mg/mL, 2 mg/mL, and 4.5 mg/mL were tested, with 0.75 mg/mL proving optimal and thus used for all subsequent experiments. The collagen I hydrogel was prepared by mixing the components on ice with 1% BSA in DPBS pre-coated wide-bore pipette tips. The pH of the hydrogel was tested with colorimetric pH strips and adjusted to pH 7.5, with 10M sodium hydroxide (Sigma-Aldrich, S5881). The hydrogel was freshly prepared for each experiment and kept on ice until combined with organoids.

### Floating gastric single- and multi-regional assembloid culture

On day 7 after seeding as single cells, gastric organoids were incubated in Cell Recovery Solution (Corning 354253) for 45 minutes on ice to dissolve the MGFR. Whole organoids were transferred to a 15 mL tube and washed twice in 10 mL of ADMEM/F12+++ and then centrifuged at 200 g for 5 minutes at 4°C. Organoids were then resuspended in 200 μL of collagen I pre-gel to be seeded and then plated in an ultra-low attachment 24-well tissue culture plate (Corning, 3473) in a ring shape for SRA. For MRA, fundus, body, and antrum organoids were seeded in the custom-designed plate. Specifically, two 30 μL drops of confluent fully grown organoids were used per region (total 6 drops per PDMS well). After being release from MGFR as describe above, each region was encapsulated in 40 μL of collagen I pre-gel (total 120 μL per PDMS well). After pre-coating the PDMS wells with 1% BSA in DPBS, each region was loaded into a separate 200 μL pipette and delivered into the well simultaneously to recapitulate the fundus, body, antrum order of normal stomach. The plate was then incubated at 37°C and 5% CO_2_ for 30 minutes to allow polymerization of the collagen-organoid. The gel structure was then detached from the bottom of the well by forceful addition of 500 μL gastric organoid medium. Medium was changed every 1-2 days (depending on cell replication) until conclusion of the experiment at day 10.

### Subcutaneous implantation of gastric assembloids in mice

Non-obese diabetic severe combined immunodeficient interleukin 2 receptor gamma null (NOD scid gamma, NSG, NOD.Cg-Prkdc^scid^ Il2rg^tm1Wjl^/SzJ) mice from Charles River Laboratories were used in all *in vivo* implantation experiments. Mice were group housed in individually ventilated cages in a specialised immunodeficient colony room, in in 12-hour light-dark cycles, with food and water ad libitum. Following subcutaneous implantation procedures, mice were recovered individually in new individually ventilated cages, before being returned to group housing with other mice who had undergone the same procedure. Access to standard diet and water was immediately provided following recovery. Each assembloid construct was loaded onto a disk of liquid MGFR contained in a sterile silicone O-rings and allowed to polymerise for 15 minutes at 37°C. MRA were orientated by application of a surgical clip to the fundus end of the tube prior to loading onto MGFR. After polymerisation, the construct loaded ring was returned to GO complete medium at 37°C and 5% CO_2_ until the implantation procedure (<24 hours). NSG mice were anaesthetised using 4% inhaled isoflurane in a sterile operating theatre within the UCL ICH Animal facility. Temperature was maintained using a heated operating table and mice were monitored intra-operatively by direct observation of respiratory rate and pattern and testing of pedal reflex, allowing adjustment of isoflurane concentration (maintenance concentration 1.5-2.5%). A 1x1cm area on the dorsum of the mouse was shaved and sterilised with dilute chlorhexidine before covering the mouse with a transparent sheet to aid sterility and thermoregulation. Bupivacaine 8mg/kg prior to making a 5mm longitudinal incision under 2x optical magnification surgical loupes. Subcutaneous pockets were created by blunt dissection according to the number of samples being implanted (maximum 4 per mouse). The wound was closed with interrupted subcuticular 5-0 Vicryl Rapide™ (polyglactin 910) sutures (Ethicon, VR493) and 0.1mg/kg buprenorphine was administered subcutaneously for analgesia.

### Retrieval and processing of implanted constructs

Constructs were retrieved after 4 weeks of implantation. Mice were sacrificed by cervical dislocation and constructs were dissected from the mouse and silicone O-ring under a Zeiss Discovery V20 SteREO microscope with a Zeiss CL 1500 HAL light source at 10-20x magnification. Images were acquired on the Zeiss microscope using a Zeiss Axiocam 506 colour camera. Samples for immunofluorescence were fixed in 4% paraformaldehyde (PFA; Sigma-Aldrich, 100496) for 30 minutes before storage in DPBS with 1X antibiotic-antimycotic (Gibco, 15240096) at 4°C. For molecular analyses, MRA constructs were divided with a scalpel into fundus, body, and antral regions with junctional zones discarded to avoid cross-region contamination, then minced and placed in RLT lysis buffer for RNA extraction.

### Immunofluorescence staining

Tissues, organoids and assembloids were fixed in 4% PFA prior processing. For tissue slides, blocking and permeabilization was done using 0.5% Triton X-100 in 1% BSA for 2 hours at RT and then primary antibodies were incubated in blocking-permeabilization solution for 24 hours at 4°C. Primary antibodies were removed using three 2-hour washes in 0.5% Triton X-100 at RT before overnight incubation of secondary antibodies at 4°C. Finally, unbound secondary antibodies were washed three times in 2-hour incubations of DPBS and the slides were then mounted for acquisition. For 3D organoid and assembloid whole mount staining, fixed samples were quenched with 0.1 M NH_4_Cl for 60 minutes. After 3 further washes with DPBS, SRA were cut into at least 4 pieces to facilitate multiple immunolabelling panels, while MRA were labelled and subsequently imaged as intact structures. The BABB-based clearing protocol was used ^33^. Samples underwent dehydration in a methanol (Sigma-Aldrich, 34860) series followed by incubation in 5% hydrogen peroxide in methanol for 2 hours at 4°C. Rehydration in reverse methanol series and equilibration in DPBS was done prior to transfer of samples to permeabilization solution (20% DMSO, 2.3% glycine (Sigma Aldrich, 410225), 0.2% Triton X-100 in DPBS). Samples were permeabilised overnight at 4°C and then transferred to blocking solution (10% DMSO, 6% normal donkey serum (NDS; Sigma-Aldrich, D9663), 0.2% Triton X-100 in DPBS) for 24 hours at RT. Primary antibodies were made up in 5% DMSO, 3% NDS, 0.2% Tween-20 (Sigma-Aldrich, P1379), and 0.1% heparin (Sigma-Aldrich, H3149) in DPBS and incubated with samples for 24 hours at 4°C. Primary antibodies were washed from the constructs with six 1-hour washes in 0.2% Tween-20 and 0.1% heparin in DPBS. Secondary antibodies were prepared in the same solution as primary antibodies and applied to the samples overnight at 4°C. Secondary antibodies were removed by six 1-hour washes in 0.2% Tween-20 and 0.1% heparin in DPBS until RI matching and imaging. All antibodies are listed in **Supplementary Table 4**.

### Acridine orange

Live MRA were cut longitudinally into halves. Cut samples were incubated 1h at 37°C with Acridine orange 10 μM (A1301, Thermo Fisher), with or without Histamine 100 μM (H7127, Sigma). Images were acquired with a Zeiss Axio Observer A1 microscope. Live experiment was carried out by incubating the acridine orange for 15 minutes at 37°C. The Histamine was either added or not, being the first time point of the experiment. Images were acquired in an automated way every 5 minutes for 1 hour (chamber at 37°C), with 20 mm thick Z-slices for 3D reconstruction.

### Microscopy image acquisition

Brightfield images of organoids in culture were acquired using on a Zeiss Axio Observer A1 inverted widefield microscope with a Colibri 5 LED light source and Axiocam MRm camera system. Reporter cell lines were imaged in culture using the same system. Sectioned samples were acquired on either a Zeiss Axio Observer A1 microscope with a Colibri 7 LED light source and Hamamatsu Flash 4v3 camera system or on a Zeiss LSM 710 inverted single photon confocal microscope. Widefield images were acquired using short Z-stacks to allow later processing by deconvolution and Gaussian-based stack focusing for 2D projection. Whole mount samples were acquired on the Zeiss LSM 710 confocal microscope on a FluoroDish. Acridine orange live imaging acquisition was performed with Nikon Eclipse Ti2 inverted microscope.

### Image processing and analysis

Brightfield images were processed using Zeiss Zen Pro. Tissue sections were processed using batch deconvolution in Huygens Essential (Scientific Volume Imaging, compute engine 21.10.0p0.64b). Whole mount image datasets were imported to Fiji for processing. Colour and contrast were minimally adjusted. Stacks were represented as Z-projections using maximum intensity projection (MIP) using all or selected slices, including single slices where this allowed optimal visualisation of the data. Whole mount image datasets of SRA and MRA were exported from Zeiss Zen Pro to IMARIS x64 (Bitplane, version 8.0.2, build 36053) for 3D and 3D stereo visualisation using the Surpass 3D view. Rendering in 3D was done using transparent (MIP) 3D mode and sections performed with the OrthoSlicer and Clipping tools. Single and multi-channel images were exported after optimisation of brightness and contrast in TIFF file format. Videos were made using the inbuilt IMARIS point-to-point animation tool and then exported directly in MP4 file format.

### Transmission electron microscopy

Gastric assembloid samples were fixed in 2.5% glutaraldehyde in 0.1 M sodium cacodylate buffer followed by secondary fixation in 1.0% osmium tetroxide. Tissues were dehydrated in graded ethanol, transferred to a transitional fluid, propylene oxide, and then infiltrated and embedded in Agar 100 epoxy resin. Polymerisation was at 60°C for 48 hours. 90 nanometre ultrathin sections were cut using a Diatome diamond knife on a Leica Ultracut UC7 ultramicrotome. Sections were picked up on Athene 300 mesh copper grids and stained with 70% alcoholic uranyl acetate and Reynold’s lead citrate for contrast. The samples were examined in a JEOL 1400 transmission electron microscope and images recorded using AMT XR80 digital camera and software. Fundus, body, and antrum regions were examined from each patient culture. Cells showing ultrastructural features were recorded at higher magnifications for further evaluation.

### Iodinated microfocus computed tomography (IMF-mCT)

Implanted SRA were iodinated by immersion overnight in 1.25% potassium triiodide (Chemondis, II0181) diluted 1:2 in 10% formalin (Sigma-Aldrich, HT501128). After washing with distilled water, samples wrapped in laboratory film and mounted in nutrient agar (Thermo Scientific, BO0336B) in a 1.5mL tube to separate it from the container during scanning. Micro-CT images of the specimen were acquired with a MedX mCT scanner (Nikon Metrology) using a molybdenum target. X-ray energy was 90 kV with a beam current of 89 μA. Exposure time was 708 msec, at 4 frames per projection, with the number of projections optimized for the size of the specimen. The resulting total average scan time was 2 hours with a voxel size of 4.33 µm. Projection images were reconstructed with modified Feldkamp filtered back-projection algorithms with proprietary software (CTPro3D, Nikon Metrology) and post processed with VGStudio MAX (Volume Graphics GmbH, Version 3.2).

### Library preparation, sequencing, alignment, and initial processing

RNA extraction was performed using RNeasy Mini Kit (Qiagen) following manufacturer’ instructions. For library preparation, total RNA quantification and integrity was confirmed using Agilent’s 4200 Tapestation (Standard Total RNA assay). For each sample, 10ng of total RNA were processed using the KAPA mRNA HyperPrep Kit (Roche p/n KK8580) according to manufacturer’s instructions. Briefly, mRNA was isolated from total RNA by use of paramagnetic Oligo dT beads to pull down poly-adenylated transcripts. The purified mRNA was fragmented using chemical hydrolysis (heat and divalent metal cation) and primed with random hexamers. Strand-specific first strand cDNA was generated using Reverse Transcriptase in the presence of Actinomycin D. This allows for RNA dependent synthesis while preventing spurious DNA-dependent synthesis. The second cDNA strand was synthesised using dUTP in place of dTTP, to mark the second strand. The resultant cDNA is then “A-tailed” at the 3’ end to prevent self-ligation and adapter dimerisation. Full length xGen adaptors (IDT), containing two unique 8bp sample specific indexes, a unique molecular identifier (N8) and a T overhang are ligated to the A-Tailed cDNA. Successfully ligated cDNA molecules were then enriched with limited cycle PCR (14 or 16 PCR cycles). The high-fidelity polymerase employed in the PCR is unable to extend through uracil. This means only the first strand cDNA is amplified for sequencing, making the library strand specific (first-strand). For sequencing, high yield, adaptor-dimer free libraries were confirmed on the Agilent TapeStation 4200 (High Sensitivity Agilent DNA 1000 assay) and a library quantification was estimated using the Qubit dsDNA HS assay (Life Technolgies). Libraries were normalised to 4nM and then an equal volume of each were pooled together. The library pool was denatured and sequenced on the NextSeq 2000 instrument (Illumina, San Diego, US) at 750pM, using a 56bp paired-read run with corresponding 8bp unique dual sample indexes and 8bp unique molecular index. For data analysis, run data were demultiplexed and converted to fastq files using Illumina’s BCL Convert Software v3.7.5.

### RNA-sequencing analysis

The identified genes were filtered out if not expressed at a count-per-million (CPM) higher than 1 in at least 2 samples of at least one condition. Data were normalized using the trimmed mean of M-values normalization method (TMM) implemented in edgeR package (version 3.40.2) ^43^. A Principal Component Analysis (PCA) was performed by R function prcomp on log2(CPM+1) data, after centering. Data visualization was performed with R package ggplot2 v. 3.4.2 ^44^. Differentially expressed genes (DEGs) were computed with edgeR, using a mixed criterion based on p-value, after false discovery rate (FDR) correction by Benjamini-Hochberg method, lower than 0.01 and absolute fold change higher than 2, unless otherwise specified. Hierarchical clustering and heat map visualization were performed using R package pheatmap v.1.0.12 with Euclidean distance and complete linkage. Functional enrichment analysis of Reactome pathways (updated in 2022) was performed by right-sided hypergeometric test using ClueGO (version 2.5.10) ^45^ and CluePedia (version 1.5.10) ^46^ within Cytoscape v. 3.10.0 environment ^47^. Proteins playing a role as ligands or receptors were selected according to Ramilowski et al. ^48^

Receptors identified as DEGs in the comparison of MRA vs. organoids or SRA in a specific subtissue, were used to subselect the corresponding ligands. The receptor-ligand pairs were further shortlisted by including only ligands that were DEGs and almost not expressed in the subtissue of interest in organoids or SRA. RNA-seq data are publicly available in GEO repository, accession number GSE247115.

### Thiol organoid barcoding in situ mass cytometry (TOB*is* MC) analysis

For cell dissociation, we optimised and adapted the published TOB*is* protocol^29,30^. Prior to barcoding, pre-treated and fixed MRA containing RFP and GFP reporter cell lines were cut with a scalpel into several pieces. Each piece was assigned a different barcode to preserve the spatial orientation in the analysed samples. Control organoids in MGFR culture were assigned separate barcodes. Barcodes were premade by the Tape lab and were applied to the samples overnight at 4°C in rotation. Unbound barcodes were quenched using 1 mM glutathione (Sigma-Aldrich, G6529) in Maxpar cell staining buffer (CSB; StandardBioTools, 201068). Dissociation solution was prepared fresh for each experiment, consisting of 0.5 mg/mL dispase II (Thermo Fisher, 17105041), 1 mg/mL collagenase IV (Thermo Fisher, 17104019), and 0.2 mg/mL DNase I (Sigma-Aldrich, DN25) final working concentrations in PBS. Samples were placed in gentleMACS C-tubes (Miltenyi Biotec, 130-093237) in 5 mL of dissociation solution, and run on the gentleMACS Octo Dissociator (Miltenyi Biotec) at 37°C in 15-minute cycles for 3-5 cycles during the optimisation phase. Each cycle consisted of forward-spin 50 rpm for 1 minute, backward-spin 50 rpm for 1 minute, then forward-spin 1500 rpm for 2 seconds, backward-spin 1500 rpm for 2 seconds, forward-spin 100 rpm for 1 minute (looped x10). After dissociation, samples were centrifuged, washed twice in CSB, and filtered through a 40 µm PET filter. A sample of filtered suspension was stained with Trypan Blue (Thermo Fisher, 15250061), counted in a Countess automated cell counter (Thermo Fisher), and inspected on a brightfield microscope for cell morphology and the presence of doublets and undigested ECM. For staining using rare earth metal-conjugated antibodies, antibodies for cell type were selected from an optimised panel. BSA- and azide-free formulations of the antibodies, using the same clone as in immunofluorescence where available, were custom conjugated to rare earth metals using Maxpar X8 polymers (StandardBioTools, 201300). Antibodies used in TOB*is* MC experiments are shown in **Supplementary Table 5**. Antibodies against extra-cellular epitopes were made up in 50 µL CSB at optimised concentrations and incubated with cells for 30 minutes at RT on a rocker. After washing with CSB, cells were permeabilised with 0.1% Triton X-100 in PBS for 30 minutes at RT on a rocker, washed with CSB again, then further permeabilised with 50% methanol in PBS for 10 minutes on ice, and washed with CSB. Antibodies against intra-cellular epitopes were made up in 50 µL CSB and applied to the cells for 30 minutes at RT on a rocker. Next, stained cells were re-fixed with 1.6% formaldehyde in PBS (Pierce, 28906) for 10 minutes at RT. After washing with CSB, DNA intercalation was performed using 125 nM Cell-ID Intercalator-Ir (StandardBioTools, 201192A) in Maxpar Fix and Perm Buffer (StandardBioTools, 201067) overnight at 4°C in which cells were stored until run on the CyTOF XT™ (StandardBioTools). For mass cytometry data acquisition, stained cells were washed with CSB and then resuspended in MaxPar CAS+ solution (StandardBioTools, 201244) at ∼0.5-1.0 x 10^6^ cells per mL. EQ6 beads (StandardBioTools, 201245) were added at 1:5 volumetric ratio and EDTA to a final concentration of 2 mM. Cells were loaded into the CyTOF XT™ and events were acquired using StandardBioTools CyTOF® software aiming for 100-400 events per second. Raw data was processed using StandardBioTools CyTOF® software to remove EQ beads and normalise signals prior to exporting data as FCS files.

After acquisition, the processed file was debarcoded (https://github.com/zunderlab/single-cell-debarcoder) and uploaded onto the Cytobank platform for further analysis. Single cells were gated for Gaussian parameters, Cisplatin^low^ and DNA^high^ to remove debris, doublets, and any other non-cellular resembling matter from the data. UMAP analysis was performed using the default settings in Cytobank (https://www.cytobank.org/). The individual debarcoded files were downloaded from Cytobank as text files and using the Mass Cytometry data analysis pipeline, CyGNAL (https://github.com/TAPE-Lab/CyGNAL), they were pre-processed and the Earth Mover’s Distance (EMD) scores were computed. Once computed, heatmaps and PCA plots were generated and compiled using OmniGraffle professional.

### WES (Simple Western™ Automated Western Blot Systems)

Protein extracts were prepared in RIPA buffer (R0278, Sigma) with protease inhibitor 1X (ab270055, Abcam) and quantified with Pierce BCA Protein Assay Kit (23227, ThermoFisher). Protein extracts were diluted at 0.3mg/mL with Antibody diluent 2 (042-203, Bio-techne) and used for Simple Western analysis, using Wes machine (004-600, Bio-techne). 12-230 kDa separation module (004-600, Bio-techne) was used, combined with the primary antibody ATP4B (MA3-923, ThermoFisher) used at 1:25 dilution, and the anti-mouse detection module (DM-002, Bio-techne). Sample normalization was done using the total protein detection module for chemiluminescence (DM-TP01, Bio-techne). Results were visualized using the Compass for Simple Western Software.

### Real-time PCR

RNA reverse transcription was performed using the High-Capacity cDNA Reverse Transcription Kit (Thermo Fisher), according to manufacturer’s instructions. Reverse transcription was done using the T100 thermal cycler (Bio-Rad) the qRT-PCR was performed with custom designed Thermo Fisher probes (Supplemenatary table 6) using Universal Sybr Green Master Rox (Merck Life Sciences, 4913850001) according to manufacturer’s instructions. Reactions were performed on QuantStudio™ 5 (Thermo Fisher) and results were analysed with QuantStudio Design & Analysis Software (v1.5.2, Thermo Fisher). GAPDH expression was used to normalize Ct values for gene expression, and data were shown as relative fold change to controls (Antrum organoids), using ΔΔCt method, and presented as mean ± SEM.

## Data availability

The authors declare that all data supporting the findings of this study are available within the article, its Supplementary Information, Source Data files, and online deposited data. The Gastric RNA-seq data generated in this study have been deposited in the NCBI GEO database. The following secure token has been created to allow review of record X_X_X_X while it remains in private status: X_X_X_X. The thiol organoid barcoding in situ mass cytometry (TOBis MC) data are deposited on X_X_X_X.

## Acknowledgments

This research was funded by the OAK Foundation Award W1095/OCAY-14-191, The Great Ormond Street Hospital (GOSH) Children’s Charity, and the National Institute for Health Research Great Ormond Street Hospital Biomedical Research Centre (NIHR GOSH BRC). BCJ is supported by the General Sir John Monash Foundation, Australia, and the UCL Overseas and Graduate Research Scholarships. GB is supported by the OAK Foundation and the UCL ICH PhD studentship. GC is supported by a UCL Division of Surgery and Interventional Science PhD scholarship. LT is supported by an NIHR UCL BRC-GOSH Crick Clinical Research Training Fellowship. VSWL is funded by the Francis Crick Institute, which receives its core funding from Cancer Research UK (CC2141), the UK Medical Research Council (CC2141) and the Wellcome Trust (CC2141). GGG is supported by the NIHR GOSH BRC, and the UCL Therapeutic Acceleration Support (TAS) LifeArc Fund Rare Diseases Call (TRO Award Number: 184646). PDC is supported by National Institute for Health Research Professorship and the GOSH Children’s Charity. Thanks to Dr Ania Rybak, Department of Gastroenterology, for the acquisition of endoscopic samples used in this study, and all paediatric surgeons in the Department of Specialist Neonatal and Paediatric Surgery. We thank Dr Valerie Karaluka and Pei Shi Chia for their assistance with sample collections. We thank Dale Moulding and Daniyal Jafree for support with BABB protocol optimization. The views expressed are those of the authors and not necessarily those of the NHS, the NIHR, or the Department of Health. The human embryonic and fetal material was provided by the Joint MRC/Wellcome (MR/R006237/1) Human Developmental Biology Resource (www.hdbr.org). We thank the UCL ICH Genomic facility.

## Author contributions

BCJ, GB, GGG and PDC designed the study. BCJ, GB and GGG performed the main experiments on organoid cultures. BCJ and GC worked on organoid, assembloid and tissue characterization. BCJ performed *in vivo* experiments. GB performed functional parietal analysis. LT and VSWL designed the organoid tagging and helped on *in vivo* experiments. ICS and OA performed Micro-CT analysis. RL designed the assembloid culture wells. MB and GA performed TEM experiments and analyses. KJ collected diseased samples and contributed to data analysis of the disease modelling. RA and CL performed sequencing analyses. JS and CT performed CyTOF experiments and analyses. SE and NE helped in the discussion and interpretation of results. GGG and GB wrote the manuscript. BCJ, GB, CT, CL, GGG and PDC critically discussed the data and manuscript.

## Inclusion and diversity

We support inclusive, diverse, and equitable conduct of research.

