## Supplementary material for "Human Gastric Multi-Regional Assembloids Favour Functional Parietal Maturation and Allow Modelling of Antral Foveolar Hyperplasia"

[1] Stem Cell and Regenerative Medicine Section, Great Ormond Street Institute of Child Health, University College London, London, UK
[2] Stem Cell and Cancer Biology Lab, The Francis Crick Institute, London, UK
[3] Department of Radiology, Great Ormond Street Hospital, London, UK
[4] Population, Policy and Practice Research and Teaching Department, Great Ormond Street Institute of Child Health, University College London, London, UK
[5] Department of Histopathology, Great Ormond Street Hospital, London, UK
[6] Department of Civil, Chemical, Environmental and Material Engineering, University of Bologna, Bologna, IT
[7] Cell Communication Lab, Department of Oncology, University College London Cancer Institute, London, UK
[8] Veneto Institute of Molecular Medicine (VIMM), Padova, Italy
[9] Department of Industrial Engineering, University of Padova, Padova, Italy
[10] Department of Paediatric Gastroenterology & Nutrition, Great Ormond Street Hospital, London, UK
[11] Department of Specialist Neonatal and Paediatric Surgery, Great Ormond Street Hospital, London, UK
* These authors contributed equally: Brendan C Jones & Giada Benedetti
[#] These authors jointly supervised this work: Giovanni Giuseppe Giobbe & Paolo De Coppi

**Extended Data**

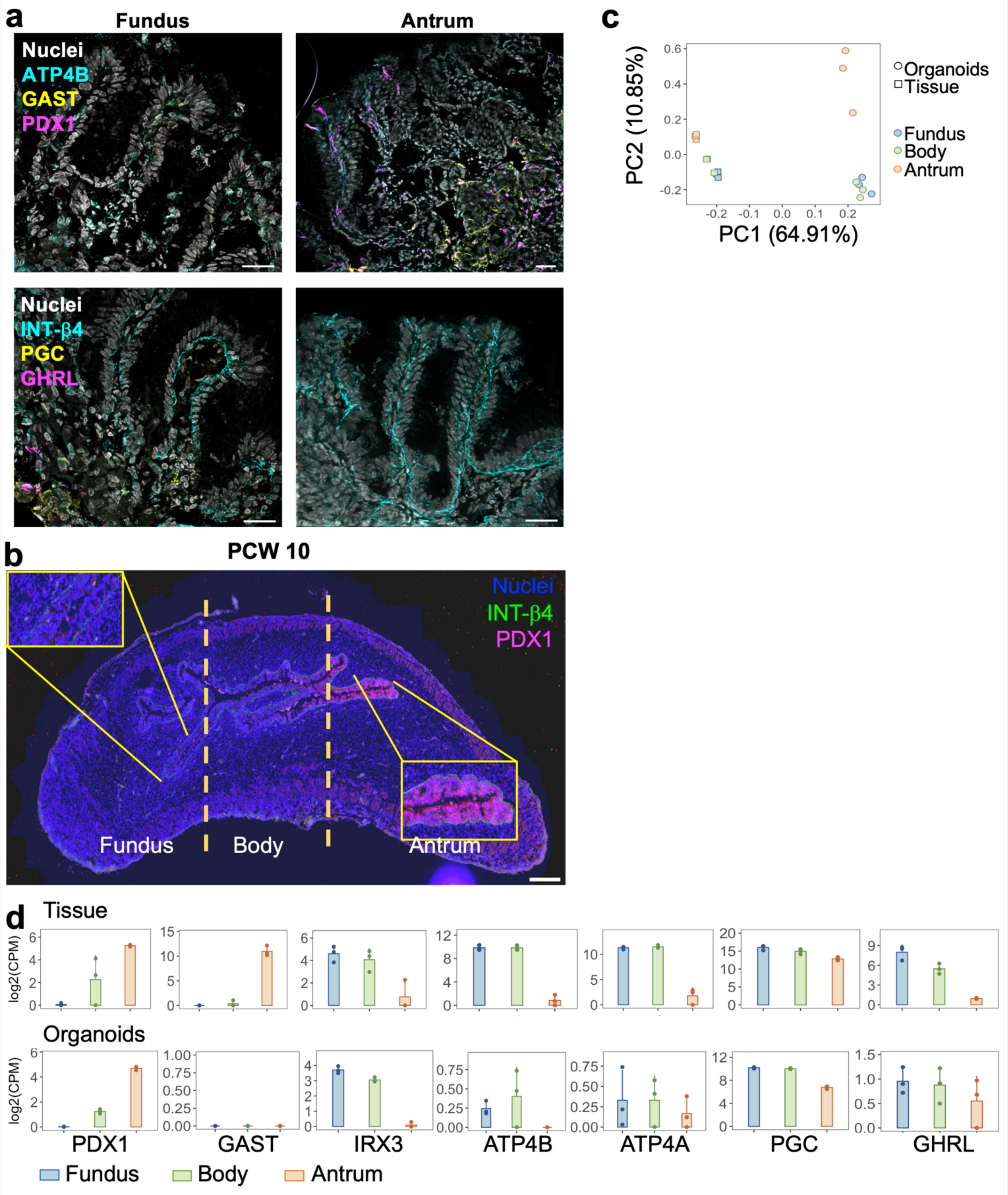

**Extended data Fig.1 | Tissue regionality and organoids characterization**

**a**, Immunofluorescence panel showing Fundus and Antrum paediatric stomach tissue sections stained for ATPase H+/K+ transporting subunit β (ATP4B) in cyan, Gastrin (Gast) in yellow, Pancreatic and Duodenal homeobox 1 (PDX1) in magenta, and nuclei in grey (Hoechst); or Integrin-β4 (INT-β4) in cyan, Pepsinogen C (PGC) in yellow, Ghrelin (GHRL) in magenta, and nuclei in grey (Hoechst). Scale bar 25 µm. **b**, Immunofluorescence panel showing foetal stomach (PCW10) tissue sections stained for Integrin-β4 (INT-β4) in green, Pancreatic and Duodenal homeobox 1 (PDX1) in magenta, and nuclei in grey (Hoechst). Scale bar 250 µm. **c**, Principal component analysis (PCA) of RNA-sequencing samples from Fundus (blue), Body (green), and Antrum (red) tissue (circles) versus organoids (squares). (n=3 for biological replicates). **d**, Expression of typical gastric markers in tissue and organoids in Fundus (blue), Body (green), and Antrum (red). Circles indicate single data points, mean ± SD (n = 3 for biological replicates) CPM: counts per million.

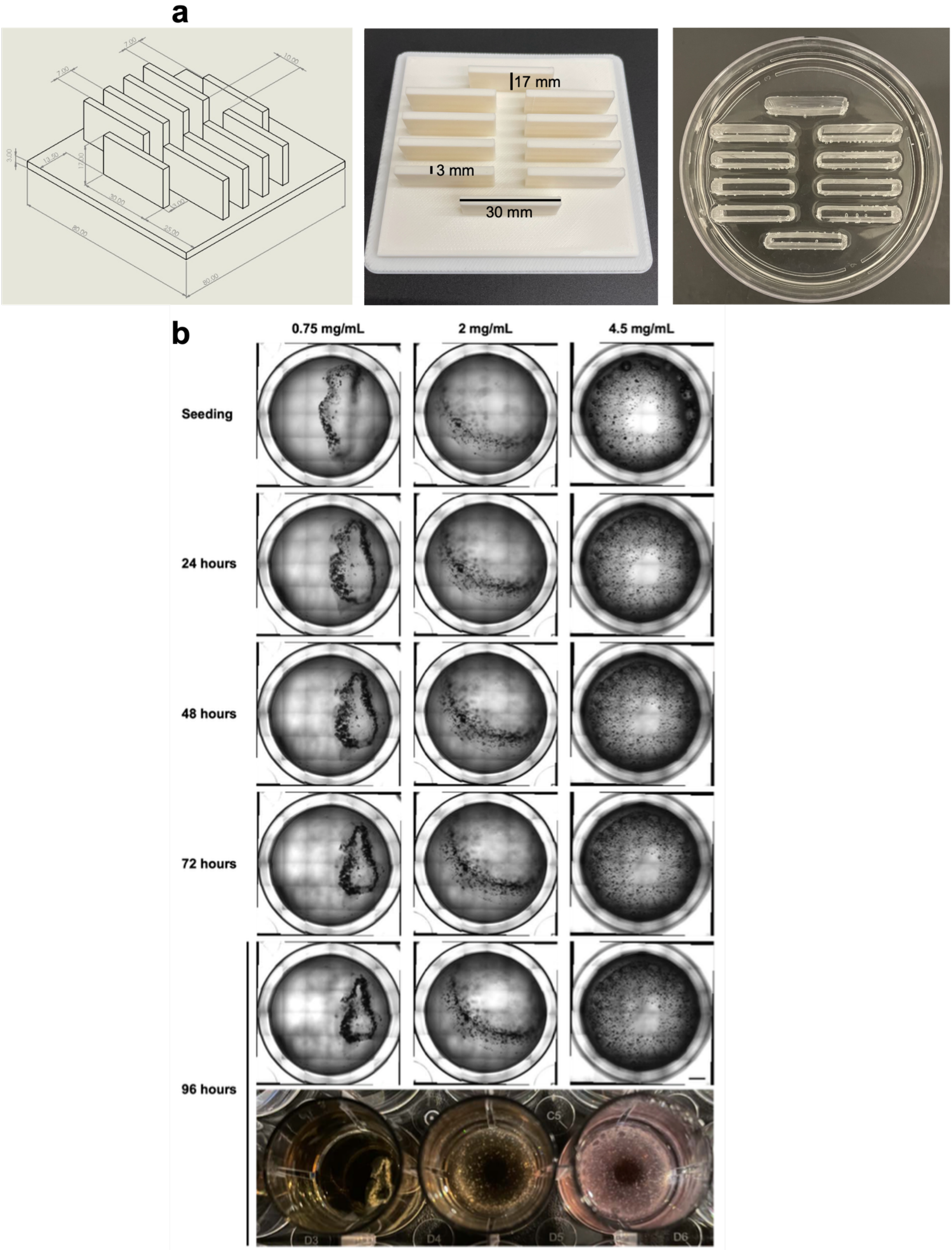

**Extended data Fig.2 | Assembloid protocol generation set-up**

**a**, Costumed-designed well for assembloid seeding: mould design (left), 3D printed mould (central), PDMS customized-well (right). **b**, SRA formation at various concentrations of collagen I hydrogel (0.75, 2, or 4.5 mg/ml). Scale bar 2 mm.

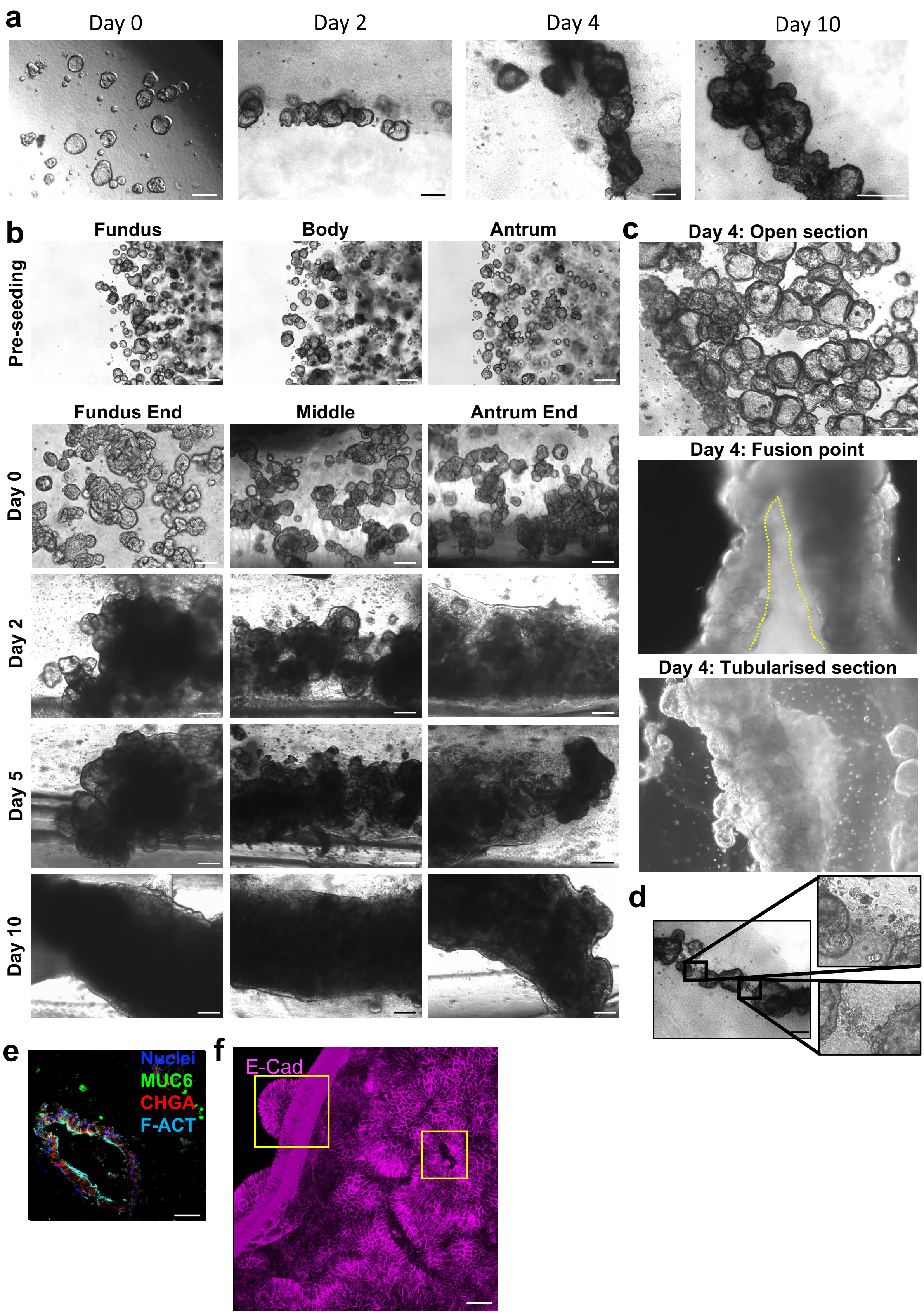

**Extended data Fig.3 | Assembloid generation**

**a**, Morphology of SRA formation when seeded at a density of 6000 organoids per mL over 10 days in floating collagen I hydrogel culture. Scale bar 200 μm. **b**, Bright field images of MRA formation at Fundus, Body, and Antrum over 10 days in floating collagen I hydrogel culture. Scale bar 200 μm. **c**, Bright field images of different parts of SRA at day 4 in culture: open section (upper panel), V-shaped fusion point marked by dotted yellow line (middle panel), tabularised section (lower panel). Scale bar 200 μm. **d**, Bright field images of different parts of SRA at day 4 in culture showing syncytium formation. Scale bar 200 μm. **e**, Immunofluorescence of sectioned gastric SRA showing mucin 6 (MUC6) in green, chromogranin A (CHGA) in red, F-actin (F-ACT) in cyan, and nuclei in blue (Hoechst). Scale bar 50 μm. **f**, Whole mount immunofluorescence of gastric SRA E-cadherin (E-Cad) in magenta (scale bar 50 μm).

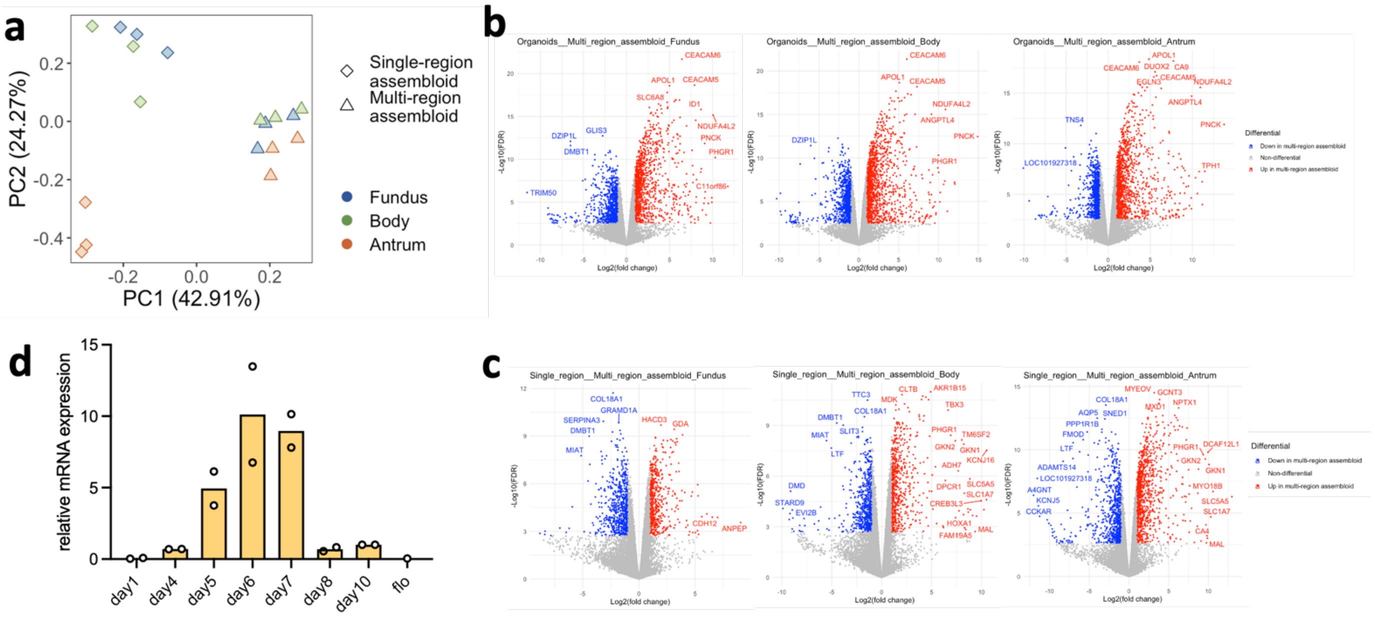

**Extended data Fig. 4 | Transcriptomics analysis**

**a**, Principal component analysis (PCA) of RNA-sequencing samples from Fundus (blue), Body (green), and Antrum (red) SRA (rhomboid) versus MRA (triangle). (n=3 for biological replicates). **b**, Vulcano plot of DEGs between organoids and MRA of Fundus, Body, and Antrum regions. **c**, Vulcano plot of DEGs between SRA and MRA of Fundus, Body, and Antrum regions. **d**, RT-qPCR for *ATP4b* whole MRA at progressive days during MRA formation, normalized to day 10. FLO (foetal lung organoids) were used as negative control. Mean and single datapoints displayed (n = 2).

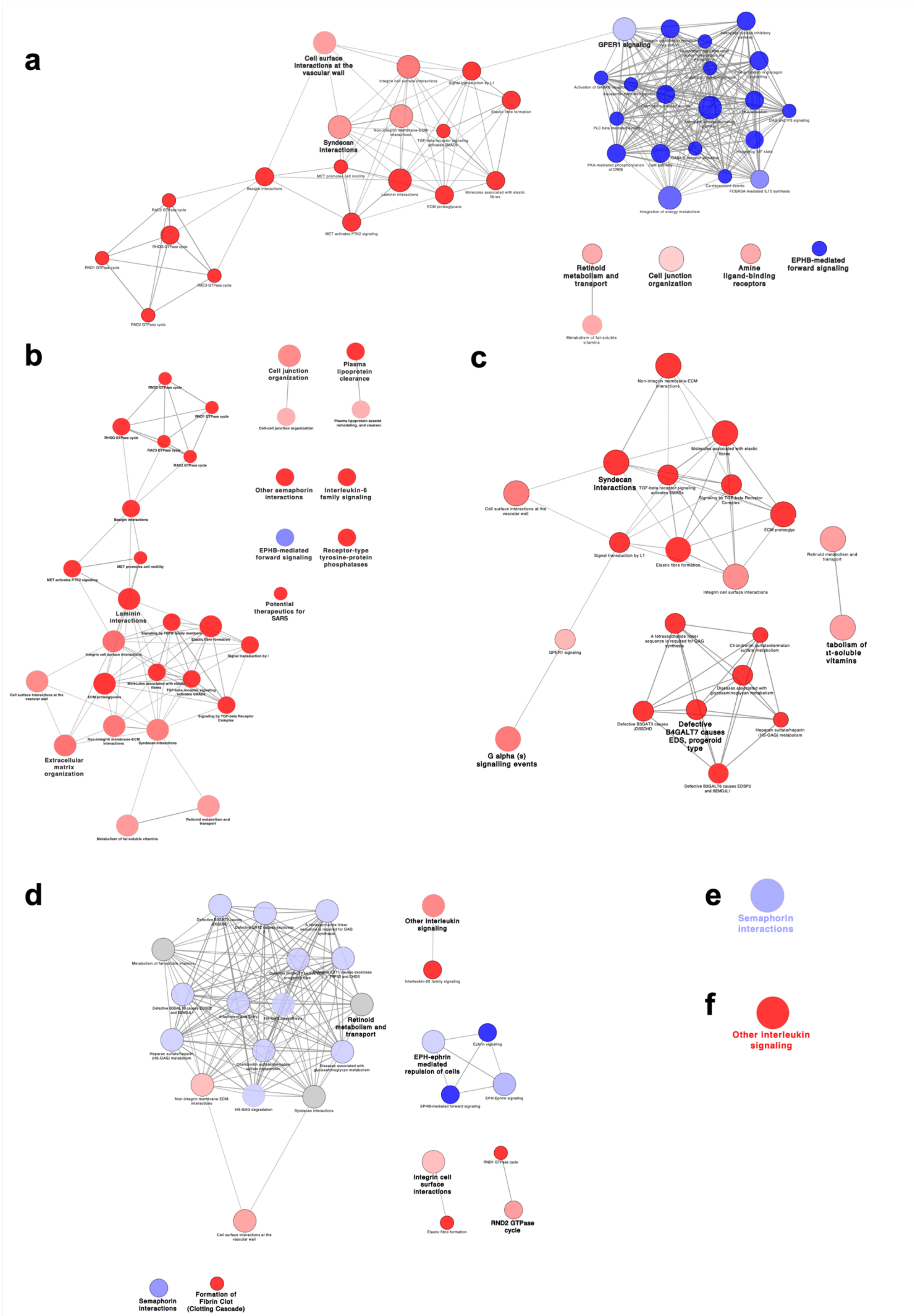

**Extended data Fig.5 | Reactome analysis**

**a**, Fundus receptor DEG reactome (organoid vs MRA). **b**, Body receptor DEG reactome (organoid vs MRA). **c**, Antrum receptor DEG reactome (organoid vs MRA). **d**, Fundus receptor DEG reactome (SRA vs MRA). **e**, Body receptor DEG reactome (SRA vs MRA). **f**, Antrum receptor DEG reactome (SRA vs MRA).

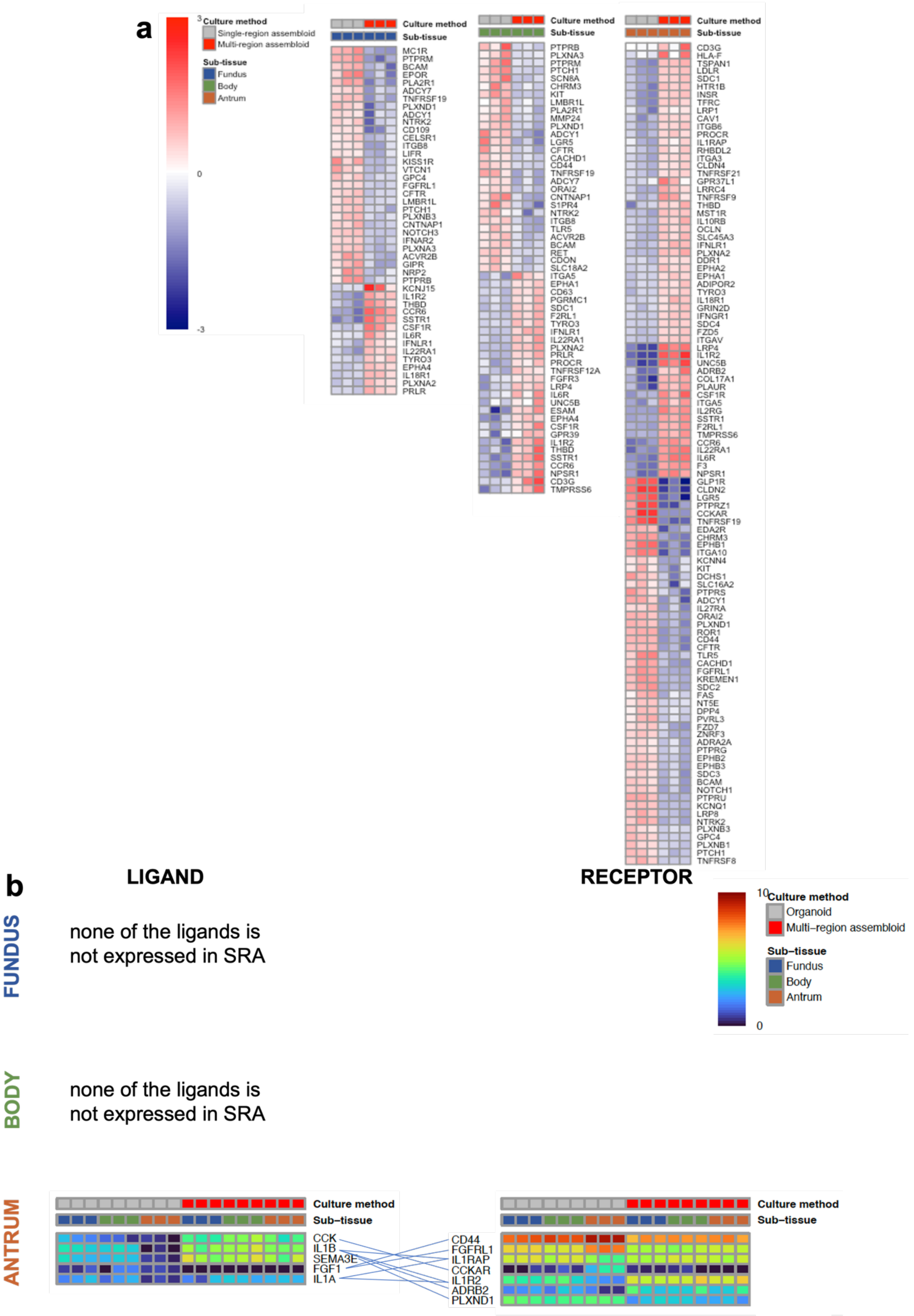

**Extended data Fig.6 | Receptor-ligand expression analysis between SRA and MRA**

**a**, Hierarchical clustering of receptors that are differentially expressed genes (DEGs) between SRA and MRA in the three region. **b**, Ligan-receptor couples expression in Fundus, Body, and Antrum in SRA vs MRA. The receptors are DEGs in the indicated subtissue between SRA and MRA. Corresponding ligands are DEGs between indicated subtissue between SRA and MRA, plus they are not expressed in the SRA of one subtissue (specified on the left).

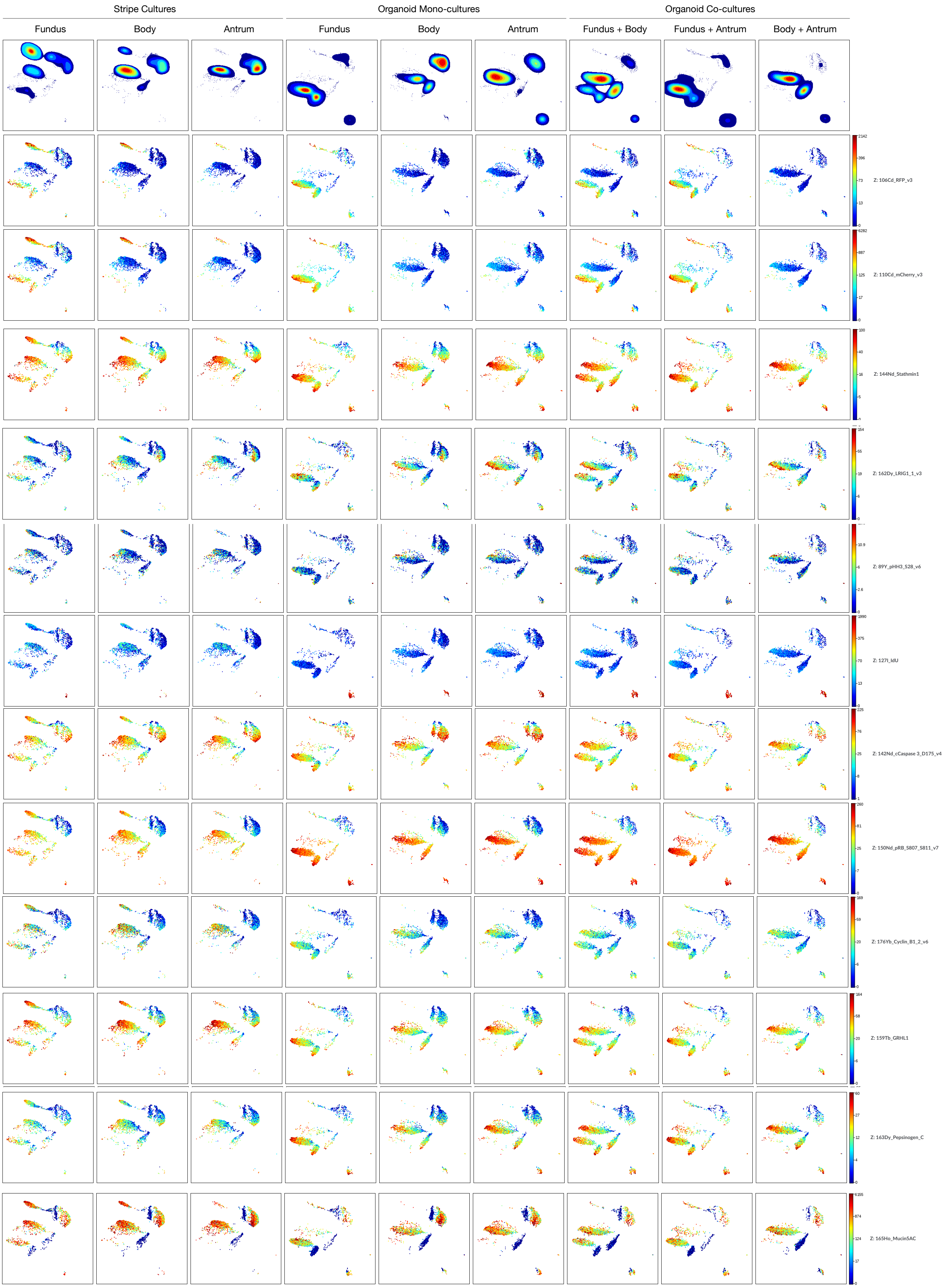

**Extended data Fig.7 |** UMAPs of the analysed markers using TOB*is* MC analysis in organoids cultures, organoids co-cultures and MRA.

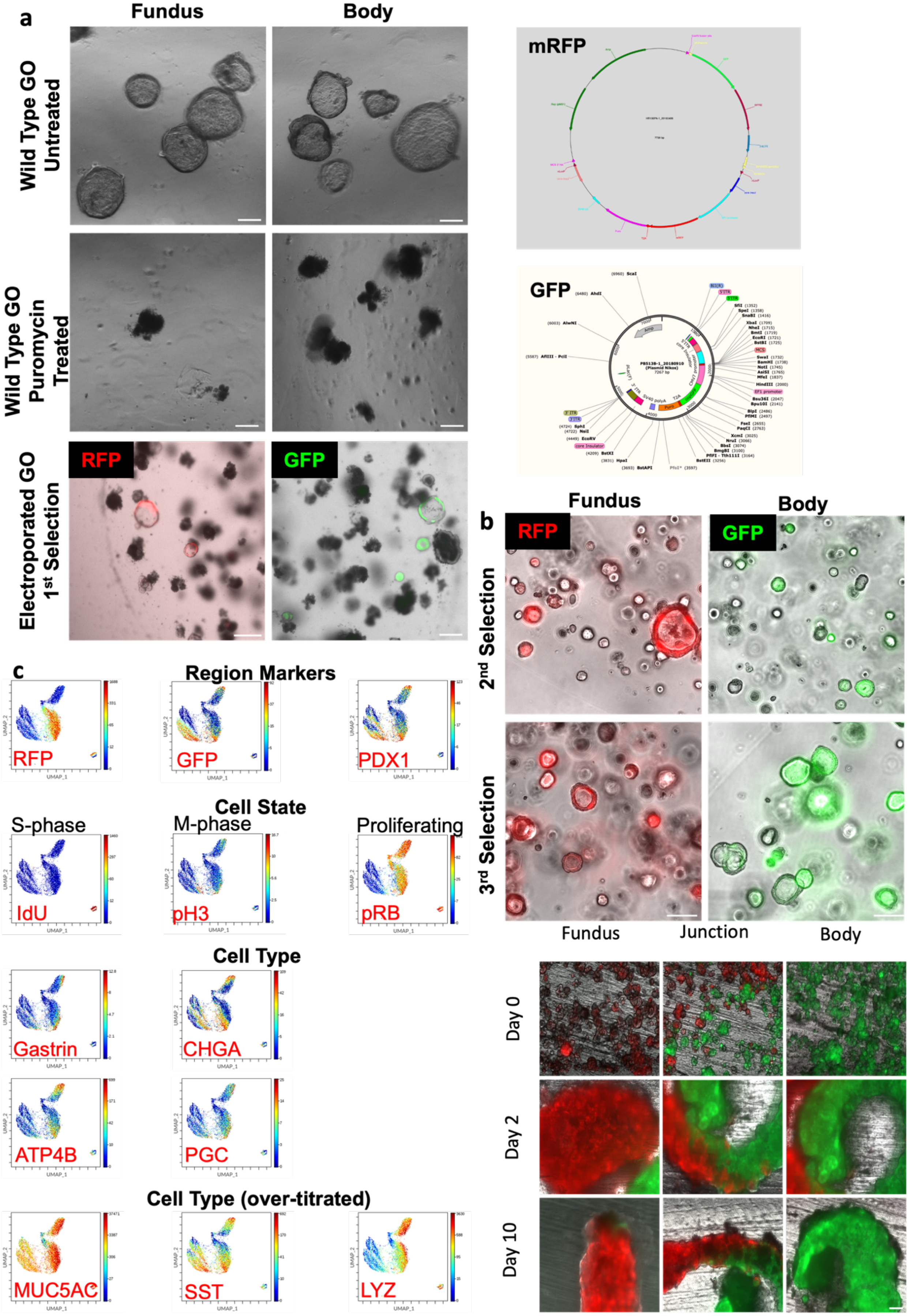

**Extended data Fig.8 | TOB*is* MC cell lines preparation and preliminary analysis**
**a**, Plasmid maps and generation of reporter RFP-fundus and GFP-body organoid lines by electroporation. Scale bars 200 µm. **b**, 2nd and 3rd selections of electroporated GOs at day 7 of passage (up). Fusion of adjacent regions in MRA (down). RFP-expressing fundus organoids, GFP-expressing body organoids. Scale bars 200 µm. **c**, Optimisation of antibody staining of MRA tubes for CyTOF.

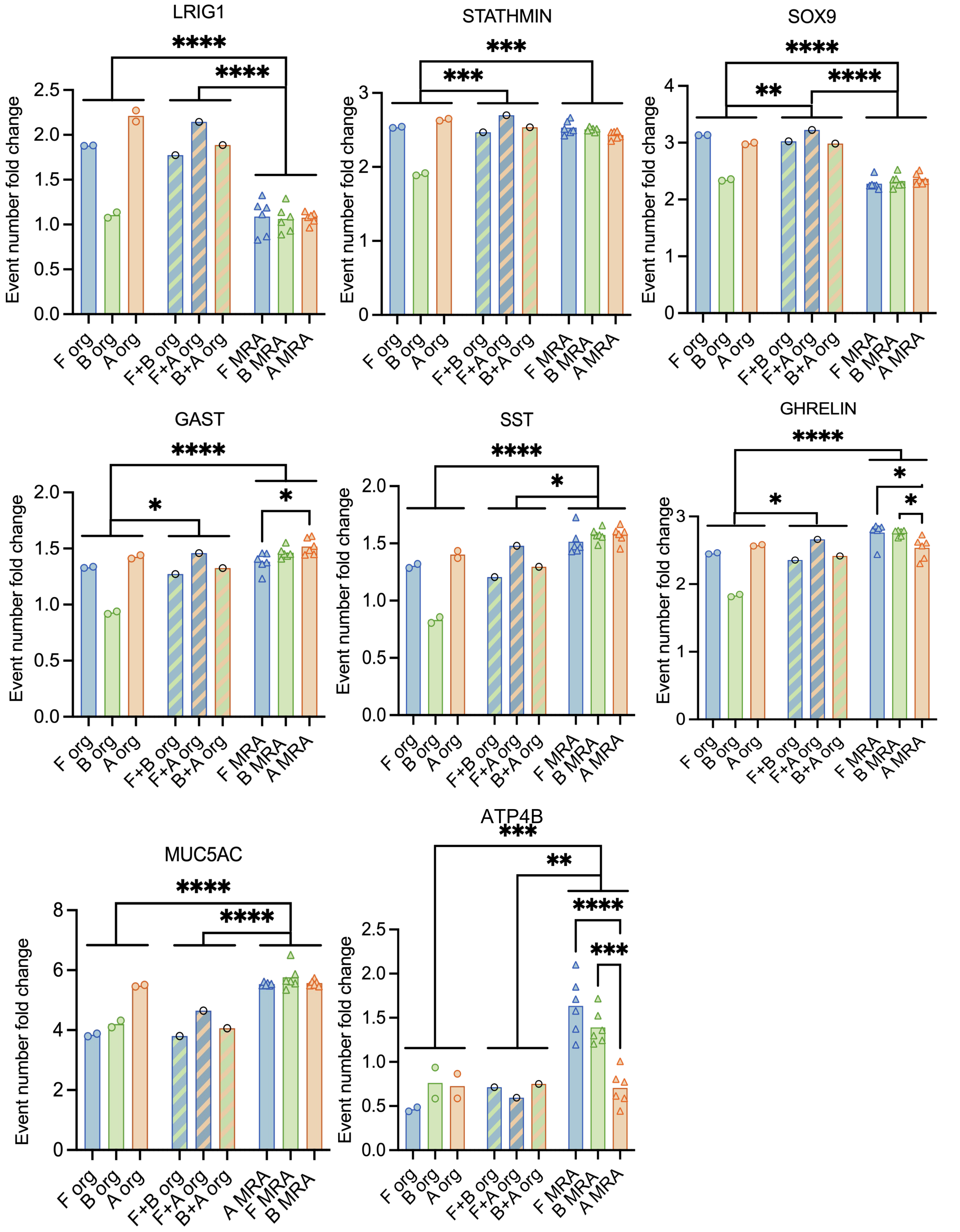

**Extended data Fig.9 | TOB*is* MC normalized reads for selected genes**
LRIG1, STATHMIN, SOX9, GAST, SST, GHRELIN, MUC5AC, ATP4B normalized reads plotted for following conditions: organoid cultures (Fundus, Body, Antrum), organoid co-cultures (F+B = Fundus and Body; F+A = Fundus and Antrum; B+A = Body and Antrum), MRA (Fundus, Body, Antrum). Mean and single data points displayed (n = 2 for organoids, n=1 for organoid co-cultures, n=6 for MRA). Two-way Anova; asterisks are graphical representation of the values of the adjusted p=value (*<0.05, **<0.01, ***<0.001, ****<0.0001).

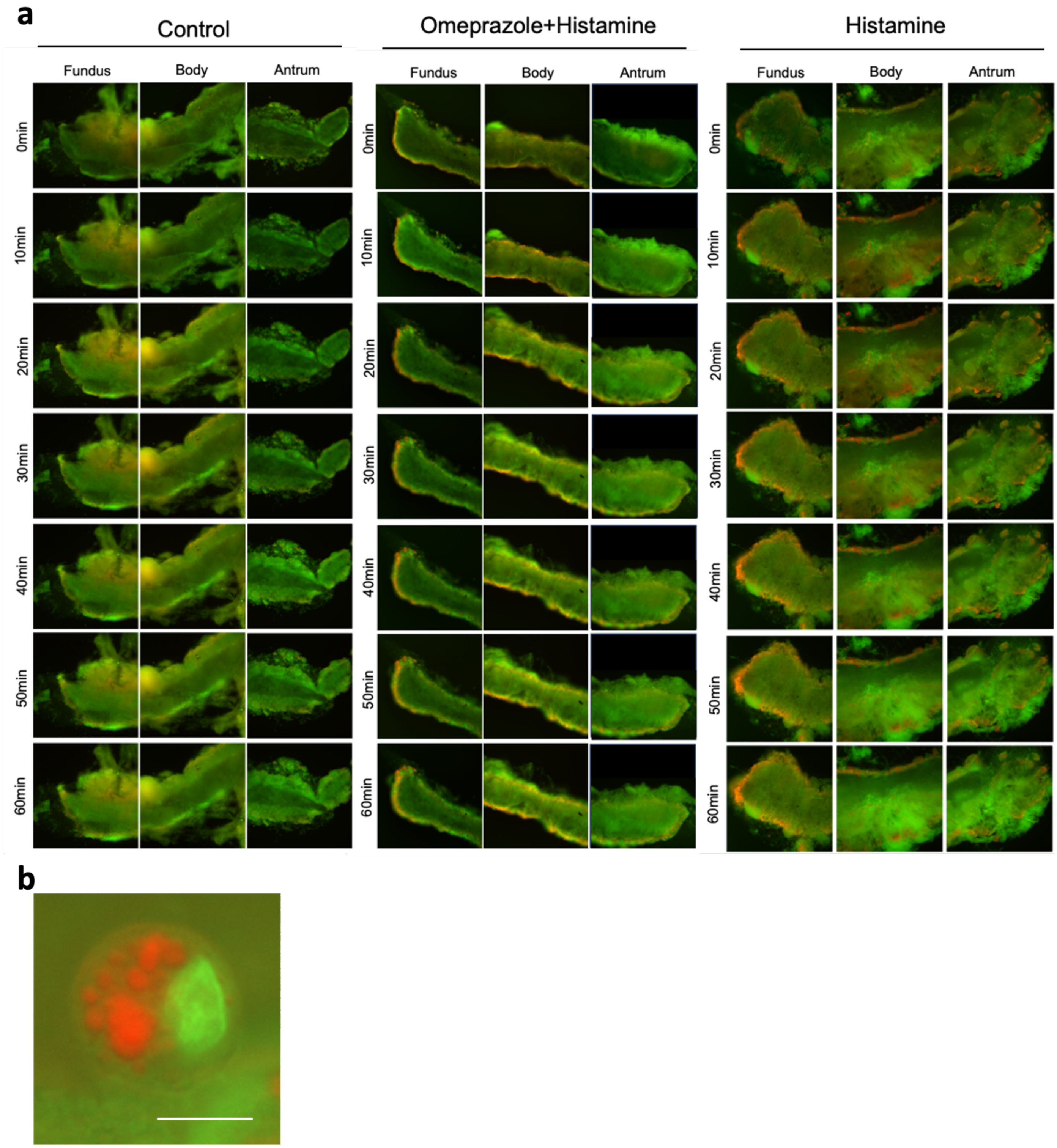

**Extended data Fig.10 | Acridine orange MRA analysis**

**a**, Timelapse frames of Acridine orange staining of Fundus, Body, and Antrum in MRA, Control, Histamine treatment (100 µM), Histamine (100 µM) + Omeprazole (100 µM). Images acquired every 10mins. **b**, 40x magnification of single cell after 1h histamine stimulation stained with acridine orange, displaying canalicular-like AO accumulation.

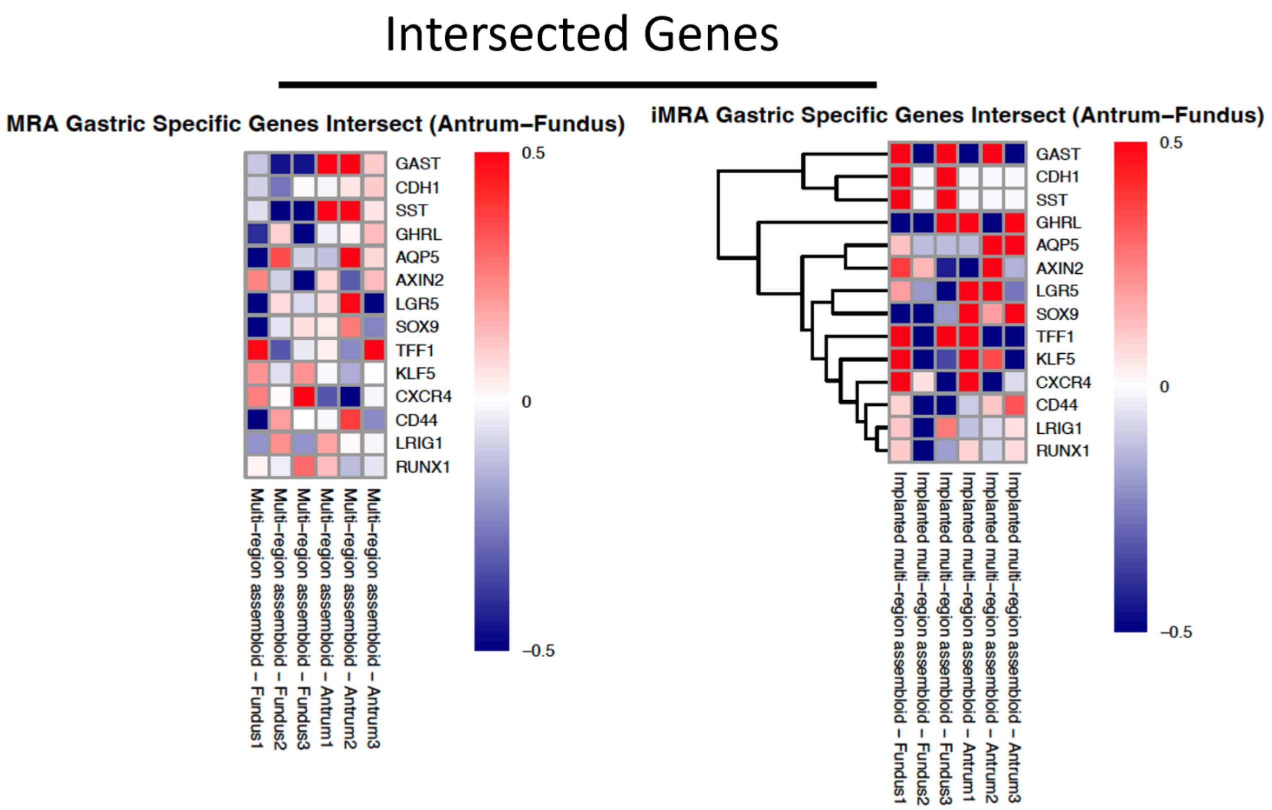

**Extended data Fig.11 |** Heatmaps of gastric specific genes between Antrum and Fundus in MRA or in iMRA. (n=3 for biological replicates).

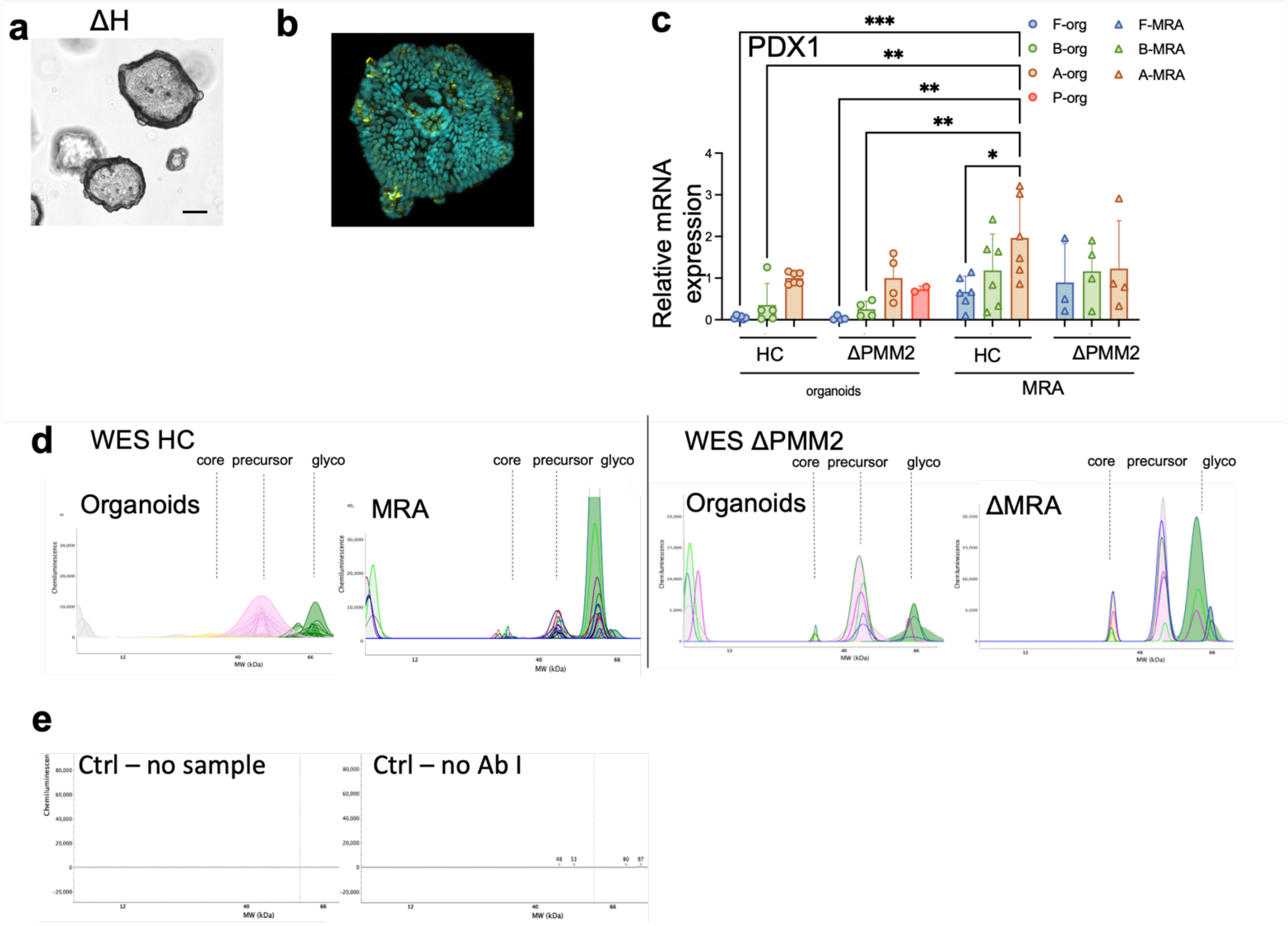

**Extended data Fig.12 | Disease modelling**

**a**, Bright field image of representative organoid line for hyperplastic region from patient affected by mutation (ΔPMM2), showing the formation of spherical organoids with visible buddings within 7 days starting from single cells. Scale bar 100 µm. **b**, Whole mount immunostaining of ΔH. **c**, RT-qPCR for Pdx1, Organoids vs MRA, HC vs ΔPMM2. (HC organoids 5 replicates, 3 patients; ΔPMM2 organoids 4 replicates, 2 patients – polyp present only in duplicate for 1 patient; HC MRA 6 replicates, 3 patients; ΔMRA 4 replicates, 2 patients. Asterisks are graphical representation of the values of the adjusted p-value from the two-way Anova (* < 0.05, ** < 0.01, *** < 0.001). **d**, Collective graphs of the peaks identified by the WES analysis (Compass Software) in Organoids vs MRA of HC on the left (n=2 per 3 patients organoids, n=3 per 3 patients MRA) and ΔPMM2on the right (n=2 technical, 1 patient only): core protein = yellow area; precursor = pink area; glycosylated protein = green area. **e**, WES control run.

**Supplementary Tables**

**Supplementary Table 1.** Gastric samples list.

| Sample | Stage | Age | Region | Use |
|---|---|---|---|---|
| 1 | Early fetal | PCW 10 | Whole | Tissue Immunostaining |
| 2 | Pediatric | 4 yo | Antrum | Tissue Immunostaining<br>Expansion<br>Sequencing<br>Organoids Immunostaining<br>Assembloid generation<br>Assembloid Immunostaining<br>CyTOF<br>In vivo implantation |
| 3 | Pediatric | 4 yo | Body | Tissue Immunostaining<br>Expansion<br>Sequencing<br>Organoids Immunostaining<br>Assembloid generation<br>Assembloid Immunostaining<br>In vivo implantation |
| 4 | Pediatric | 4 yo | Body_GFP | Expansion<br>Assembloid generation<br>CyTOF<br>In vivo implantation<br>Sequencing |
| 5 | Pediatric | 4 yo | Fundus | Tissue Immunostaining<br>Expansion<br>Sequencing<br>Organoids Immunostaining<br>Assembloid generation<br>Assembloid Immunostaining<br>In vivo implantation |
| 6 | Pediatric | 4 yo | Fundus_FRP | Expansion<br>Assembloid generation<br>CyTOF<br>In vivo implantation<br>Sequencing |
| 7 | Pediatric | 7 yo | Antrum | Tissue Immunostaining<br>Expansion<br>Organoids immunostaining<br>Assembloid generation<br>WES analysis<br>Sequencing<br>RT-qPCR |
| 8 | Pediatric | 7 yo | Body | Tissue Immunostaining<br>Expansion<br>Organoids immunostaining<br>Assembloid generation<br>WES analysis<br>Sequencing<br>RT-qPCR |
| 9 | Pediatric | 7 yo | Fundus | Tissue Immunostaining<br>Expansion<br>Organoids immunostaining<br>Assembloid generation<br>WES analysis<br>Sequencing<br>RT-qPCR |
| 10 | Pediatric | 11 yo | Antrum | Expansion<br>Organoid immunostaining |

| | | | | |
|---|---|---|---|---|
| | | | | Assembloid generation<br>WES analysis<br>AO staining<br>TEM analysis<br>Sequencing<br>RT-qPCR |
| 11 | Pediatric | 11 yo | Body | Expansion<br>Organoid immunostaining<br>Assembloid generation<br>WES analysis<br>AO staining<br>TEM analysis<br>Sequencing<br>RT-qPCR |
| 12 | Pediatric | 11 yo | Fundus | Expansion<br>Organoid immunostaining<br>Assembloid generation<br>WES analysis<br>AO staining<br>TEM analysis<br>Sequencing<br>RT-qPCR |
| 13 | Pediatric | 6mo | Antrum | Expansion<br>Organoid immunostaining<br>Assembloid generation<br>WES analysis<br>AO staining<br>Sequencing<br>RT-qPCR |
| 14 | Pediatric | 6mo | Body | Expansion<br>Organoid immunostaining<br>Assembloid generation<br>WES analysis<br>AO staining<br>Sequencing<br>RT-qPCR |
| 15 | Pediatric | 6mo | Fundus | Expansion<br>Organoid immunostaining<br>Assembloid generation<br>WES analysis<br>AO staining<br>Sequencing<br>RT-qPCR |
| 16 | Pediatric | 8yo | Antrum | Expansion<br>Organoid immunostaining<br>Assembloid generation<br>WES analysis<br>AO staining<br>RT-qPCR<br>TEM analysis |
| 17 | Pediatric | 8yo | Body | Expansion<br>Organoid immunostaining<br>Assembloid generation<br>WES analysis<br>AO staining<br>RT-qPCR<br>TEM analysis |
| 18 | Pediatric | 8yo | Fundus | Expansion<br>Organoid immunostaining<br>Assembloid generation |

| | | | | |
|---|---|---|---|---|
| | | | | WES analysis<br>AO staining<br>RT-qPCR<br>TEM analysis |
| 19 | Pediatric | 8yo | Hyperplastic region | Expansion<br>Organoid immunostaining<br>Assembloid generation<br>WES analysis<br>AO staining<br>RT-qPCR<br>TEM analysis |
| 20 | Pediatric | 8yo | Antrum | Expansion<br>Organoid immunostaining<br>Assembloid generation<br>WES analysis<br>AO staining<br>RT-qPCR<br>TEM analysis |
| 21 | Pediatric | 8yo | Body | Expansion<br>Organoid immunostaining<br>Assembloid generation<br>WES analysis<br>AO staining<br>RT-qPCR<br>TEM analysis |
| 22 | Pediatric | 8yo | Fundus | Expansion<br>Organoid immunostaining<br>Assembloid generation<br>WES analysis<br>AO staining<br>RT-qPCR<br>TEM analysis |

**Supplementary Table 2.** Gastric organoids medium composition

| **Component** | **Stock conc.** | **Final conc.** |
|---|---|---|
| Advanced DMEM F-12 (Thermo 12634) | - | To volume |
| HEPES (Thermo 15630080) | 1 M | 10 mM |
| Glutamax (Thermo 35050061) | 100 X | 2 mM |
| Pen/Strep (Thermo 15140122) | 100 % | 1 % |
| Primocin (Thermo Fisher NC9392943) | 50 mg/mL | 100 µg/mL |
| B-27 supplement minus vitamin A (Thermo 12587010) | 50 X | 1 X |
| n-acetylcysteine (Sigma A9165) | 500 mM | 1.25 mM |
| Wnt-3A (Peprotech 315-20) | 50 µg/mL | 100 ng/mL |
| R-spondin 1 (Peprotech 120-38) | 100 µg/mL | 500 ng/mL |
| Noggin (Peprotech 120-10C) | 100 µg/mL | 100 ng/mL |
| Human EGF (Thermo Fisher PMG8043) | 500 µg/mL | 50 ng/mL |
| Gastrin (Sigma G9020) | 100 µM | 10 nM |
| GSK-3 inhibitor (CHIR 99021) (Tocris 4423) | 3 mM | 3 µM |
| TGFβ inhibitor (A83-01) (Sigma SML0788) | 500 µM | 5 µM |
| FGF10 (Peprotech 100-26) | 100 µg/mL | 200 ng/mL |
| ROCK inhibitor Y-27632 (Tocris 1254) | 10 mg/mL | 10 µg/mL |

**Supplementary Table 3.** Collagen I Hydrogel composition (0.75mg/mL)

| Reagent | Company | Catalogue | Stock Concentration | Final Concentration |
|---|---|---|---|---|
| MilliQ water (18.2mΩ/cm) | Merck Millipore | ZIQ7003T0 | - | To volume |
| Advanced DMEM/F-12 | Thermo Fisher Scientific | 12500062 | 10X | 1X |
| HEPES | Thermo Fisher Scientific | 15630080 | 1 M | 10 mM |
| Collagen I (rat tail) | First Link | 60-35-810 Batch: RTC8328 | 5.05 mg/mL | 0.75 mg/mL |

**Supplementary Table 4.** Immumostaining Antibody List

| Antibody | Code | Brand | Host | Dilution |
|---|---|---|---|---|
| ATPase H+/K+ transporting subunit beta (ATP4B) | MA3-923 | Thermo | Mouse | 1:100 |
| Chromogranin A (CHGA) | ab15160 | Abcam | Rabbit | 1:50 |
| E-cadherin (E-CAD) | 610181 | BD bioscience | Mouse | 1:100 |
| Gastrin (GAST) | ab223501 | Abcam | Rabbit | 1:100 |
| Ghrelin (GHRL) | NB600-813 | Novus Biology | Goat | 1:100 |
| Integrin beta-4 (INTB4) | ab110167 | Abcam | Rat | 1:200 |
| Iroquois Homeobox 3 (IRX3) | Sc-166657 | Santa Cruz | Mouse | 1:100 |
| Lysozyme (LYZ) | GTX72913 | GeneTex | Rabbit | 1:100 |
| Mucin 5AC (MUC5AC) | ma5-12178 | Invitrogen | Mouse | 1:100 |
| Mucin 6 (MUC6) | ab216017 | Abcam | Mouse | 1:100 |
| Pancreatic and duodenal homeobox 1 (PDX1) | AF2419 | R&D SYSTEMS | Goat | 1:100 |
| Pepsinogen (PGC) | HPA031717 | Atlas | Rabbit | 1:100 |
| Somatostatin (SST) | MAB2358 | R&D Systems | Rat | 1:100 |
| anti-Mouse 488 (secondary AB) | A11001 | Thermo Fisher | Goat | 1:300 |
| anti-Mouse 568 (secondary AB) | A10037 | Thermo Fisher | Goat | 1:300 |
| anti-Mouse 647 (secondary AB) | A31571 | Thermo Fisher | Donkey | 1:300 |
| anti-Rabbit 568 (secondary AB) | A11042 | Thermo Fisher | Donkey | 1:300 |
| Anti-Rat 488 (secondary AB) | A48269 | Thermo Fisher | Donkey | 1:300 |
| Anti-rabbit 647 (secondary AB) | A-21244 | Thermo Fisher | Goat | 1:300 |
| Anti-Goat 633 (secondary AB) | A-21082 | Thermo Fisher | Donkey | 1:300 |
| anti-Rat 594 (secondary AB) | A11007 | Thermo Fisher | Goat | 1:300 |
| Phalloidin 647 | A22287 | Thermo Fisher | --- | 1:500 |
| Hoechst 33342 (nuclear staining) | H1399 | Thermo Fisher | --- | 10 μg/mL |

**Supplementary Table 5.** TOB*is* MC Antibody List

| Antibody | Code | Brand | Metal |
|---|---|---|---|
| RFP | 200-301-379 | eBiosciences | Cd |
| mCHERRY | M11217 | Thermofisher | Cd |
| IdU | Q-27194 | Fluidigm | IdU |
| cCaspase3 [D175] (v4) | #9661 | CST | Nd |
| Stathmin1 | NBP1-76798 | Bio-techne | Nd |
| pNDRG1 [T346] (v2) | #5482 | CST | Nd |
| Lysozyme | 5490-4110 | Biorad | Nd |
| pSRC [Y418] (v8) | 50-9034-42 | eBioscience | Nd |
| p4E-BP1 [T37/46 ] (v3) | #2855 | CST | Sm |
| pRB [S807/811] (v5) | 558389 | BD Biosciences | Nd |
| pAKT [T308] (v9) | 558275 | BD Biosciences | Sm |
| pCREB [S133] (v11) | #9198 | CST | Eu |
| SOX9 (v3) | Ab76997 | Abcam | Yb |
| pMKK3/MKK6 [S189/207] (v4) | #9231 | CST | Gd |
| pP38 [T180/Y182] (v4) | #9211 | CST | Gd |
| GHRL1 | H00051738-M01 | Abcam | Ho |
| Gastrin | ab223501 | Abcam | Dy |
| LRIG1 (V3) | AF7498 | R&D Syustems | Dy |
| Pepsinogen C | ab255826 | Abcam | Yb |
| pP120-Catenin [T310] (V2) | 558203 | BD Bioscinces | Dy |
| Mucin5AC | MA5-12178 | Abcam | Er |
| pGSK3b [S9] (S2) | #9336 | CST | Er |
| Mucin6 | ab216017 | Abcam | Er |
| pSMAD2 [S465/467] /pSMAD3 [S423/425] (v5) | #8828 | CST | Er |
| CHGA | sc-393941 | Santa Cruz | Er |
| Somatostatin | MAB2358 | Atlas | Yb |
| cPARP [D214] (2) (v3) | #5626 | CST | Yb |
| K-ATPase_ATP4B | Ab176992 | Abcam | Lu |
| Cyclin B1 | 554179 | BD Biosciences | Yb |
| pHistone H3 [S28] | 641010 | BioLegend | Y |
| PDX1 | AF2419 | R&D Systems | Yb |
| GFP (V2) | 50-6498-82 | eBiosciences | Cd |
| LGR5 | MAB82401 | R&D Systems | Sm |

**Supplementary Table 6.** qRT-CPR primer list

| Gene | Forward | Rerverse |
|---|---|---|
| *GAPDH* | CACATCGCTCAGACACCA TG | TGACGGTGCCATGGAATTT G |
| *MUC5AC* | GCTTTCTTGTGGAGGGTGTG | GATCACCATGTCCAAGCGTC |
| *ATP4B* | GACCTTCAACAATCCCCACG | TTCTCTGGAGTTCGCACACT |
| *PDX1* | TCCTACAGCACTCCACCTTG | ACTGGCATCAATTTCACGGG |
